# A Standardized Protocol to Investigate Trans- Endothelial Trafficking in Zebrafish: Nano-bio Interactions of PEG-based Nanoparticles in Live Vasculature

**DOI:** 10.1101/2025.07.23.666282

**Authors:** Ye-Wheen Lim, Craig A. Bell, Nicholas L. Fletcher, Nicholas D. Condon, Dewan T. Akhter, James Rae, Charles Ferguson, Nick Martel, James Humphries, Harriet P. Lo, Yeping Wu, Zachary H. Houston, Anne K. Lagendijk, Thomas E. Hall, Kristofer J. Thurecht, Robert G. Parton

## Abstract

Trans-endothelial transport of nanoparticles remains poorly characterized in live organisms. The zebrafish is a well-established model for direct *in vivo* imaging; however, standardized controls are not consistently applied across studies. Here, we developed a standardized protocol to assess nanoparticle trans-endothelial trafficking in live zebrafish. We tested a range of tracers and identified 2000 kDa dextran as an optimal control for standardizing microinjections and quantifying nanoparticle transport in a systematic unbiased manner. As validation, we identified the early physiological trans-endothelial transport pathways in zebrafish embryos using 40 kDa dextran as a nanoparticle surrogate. Using our workflow, we assessed the extravasation of 3, 7, 32 and 47 nm polyethylene glycol (PEG)-based nanoparticles. Using cAMP stimulation to restrict paracellular routes, and inhibitors to impede dynamin-dependent endocytosis, we demonstrated that PEG-based hyperbranched polymer (HBP) nanoparticles of 3 to 7 nm traverse the endothelial barrier via paracellular routes and 32 and 47 nm PEG micelles adopt dynamin-dependent endocytic trafficking over paracellular transport. We characterized the emergence of caveolae in the vasculature up to 17 days post-fertilization (dpf) using a *cavin1b* knock-in zebrafish line. Using *cavin1a/cavin1b* double knockout (DKO) zebrafish, and tumor-bearing Cavin-1 null mice, we showed that caveolae do not contribute to the transvascular transport of these PEG-based nanoparticles. This work highlights the rigour of the standardized protocol for assessing nanoparticle trans-endothelial transport in the live zebrafish, providing a systematic approach for quantification using a standardized control.

---

Intravenously administered nanoparticles must traverse multiple biological barriers to reach their target site, with the endothelial lining of the blood vasculature as one of the primary obstacles. ^1^ Elucidating the mechanisms that govern nanoparticle-cell membrane interaction at the endothelium, and nanoparticle translocation from the lumen to the extravascular space (EVS) is necessary for understanding nanoparticle delivery in biological and medical applications. ^2^ This remains a critical challenge, especially under physiological conditions. *In vitro* studies on adherent cells often fail to replicate the systemic complexity of *in vivo* environments, ^3^ while mammalian *in vivo* models face hurdles in assessing precise cell biology or high-resolution live imaging. The zebrafish, with their optical transparency and conserved vascular morphology and genetics, offer a promising vertebrate model to overcome these hurdles. ^4–6^ Pioneering studies have leveraged the zebrafish to explore the behaviour of administered nanoparticles at the cellular level, ^7^ detailing *in vivo* events of nanoparticle circulation, clearance and targeted delivery. ^8–15^ An inherent challenge remains in ensuring consistent intravenous administration of nanoparticles in a developing organism such as the zebrafish embryo. There is variability in developmental stage, particularly across clutches, ^16–18^ and variability in injection calibration, leading to inconsistency in the total concentration and volume of an intravenously injected nanoparticle solution. ^19^ Discrepancy in microinjection sites, such as the heart, posterior cardinal vein or common cardinal vein (CCV) adds to this variability. Artefactual signals of injected nanoparticles may also be detected in the blood vasculature and surrounding tissues when injections are inaccurately performed, in which compromised vessel and tissue integrity leads to unintended diffusion and transport of nanoparticles into or out of the vascular lumen. Developing a standardized framework for investigating trans-endothelial transport and biodistribution of circulating nanoparticles in the zebrafish is therefore essential.

In wild-type (WT) zebrafish with intact physiological endothelium, nanoparticles administered into the bloodstream exhibit uptake by macrophages ^7, 14, 20, 21^ and scavenger endothelial cells (SECs), ^8, 9^ with the latter characterized primarily by punctate signal in the caudal vein plexus (CVP). ^8, 9, 14^ Despite numerous reports of nanoparticle localization and activity in endothelial regions distal to the CVP, such as the intersegmental vessels (ISVs), ^7, 12, 15, 20, 22, 23^ studies on this interaction remain scarce. Nanoparticle size is a critical determinant of trans-endothelial transport mechanisms. ^2, 24, 25^ Clathrin coated vesicles, reliant on the GTPase dynamin for vesicle scission, typically form vesicles of 50 to 100 nm in diameter, extending up to 300 nm. ^2, 13^ This lumen size range aligns with the sub-200 nm size of most nanoparticles designed in recent years. ^26, 27^ Uptake of nanoparticles into endothelial cells via clathrin coated pits has been shown to occur in the zebrafish, ^13^ while recent work in mice has highlighted active trans-endothelial transport as a prominent route by which nanoparticles can travel through the tumor endothelial barrier to reach target tumor sites. ^28, 29^ This size barrier also applies to the controversial mechanism of nanoparticle cellular uptake and trans-endothelial transport via caveolae, which possess an internal diameter of typically 60 to 80 nm. ^2, 13, 30–32^ Additionally, nanoparticle size dictates their potential for transcellular transport via inter-endothelial cell openings. ^33, 34^ Tight and adherens junction openings between zebrafish endothelial cells range from approximately 3 to 6 nm and 12 to 15 nm respectively, depending on the vessel type, and development stage of the zebrafish. ^35^

The presence of polyethylene glycol (PEG) on nanoparticles can impart steric shielding, prolonging circulation time *in vivo* and reducing recognition by the immune system. ^36^ In the zebrafish, PEGylation of liposomes and polystyrene nanoparticles has been shown to reduce specific clearance by macrophages and enhance circulation time. ^8, 15, 20, 22^ These circulating PEGylated nanoparticles appeared adherent to blood vessel endothelial cells and were eventually cleared, suggesting a role for trans-endothelial transport in the interaction between PEGylated nanoparticles and endothelial cells. Understanding the mechanisms by which PEG-functionalized nanoparticles interact with the endothelial barrier *in vivo* is crucial for designing an efficient nanoparticle delivery system.

Here, we present a standardized protocol centred on the use of a tracer control, broadly applicable to studies involving the administration of nanoparticles in live zebrafish embryos and juveniles. We uncovered the trans-endothelial routes taken by PEG-based hyperbranched polymers (HBPs) and micelles with diameters ranging from 3 nm to 47 nm at a cellular level. This standardized protocol integrates genetic and chemical manipulations within its workflow, enabling the systematic elucidation of specific endocytic pathways involved in nano-bio interactions between PEG-based nanoparticles and endothelial cells *in vivo*.

## Results and Discussion

### 2000 kDa Dextran as a Tracer Control in a Standardized Protocol for Ratiometric Quantitation of Trans-endothelial Transport

To control for variability in the microinjection of a developing embryo, we explored the use of a tracer control. An optimal tracer control is able to 1. label the blood vasculature lumen of the whole embryo 2. sense perturbations in the integrity of the vasculature and surrounding tissues in the whole zebrafish 3. control for volume and concentration of co-injected nanoparticles and 4. provide a baseline signal, allowing standardized quantitation. This tracer should possess a high and detectable signal for imaging and analysis. We screened a range of fluid-phase marker dextran of varying sizes in 2 dpf WT zebrafish embryos. The 2 dpf stage was selected as 2 dpf embryos exhibit developed and lumenized longitudinal axial vessels and secondary vessels, accompanied by the establishment of blood flow. ^5, 37^ We first injected dextran-fluorophore conjugates of sizes 10 kDa [>3.7 to 5.4 nm hydrodynamic diameter (Dh)], 40 kDa (>9.56 to 11.4 nm Dh), 70 kDa (10.2 to 14.8 nm Dh) and 2000 kDa (53.78 to 54.4 nm Dh). ^38–40^ Upon injection, 10 kDa dextran showed immediate diffusion throughout the embryo, indicating rapid dispersal in the whole organism (**Figure 1A**). 40 kDa and 70 kDa dextran injected embryos showed punctate signal at the blood vessel surface and signal in the EVS 30 minutes post injection (mpi) (**Figure 1B-C**). Importantly, 2000 kDa dextran signal was specifically confined within the blood vasculature at 30 mpi (**Figure 1D**). We further examined the routes of dextran transport via live imaging of a 2 dpf zebrafish embryo injected with a mixture of 40 kDa dextran-Texas Red and 2000 kDa dextran-FITC. 40 kDa and 2000 kDa dextran signal was immediately present throughout the blood vessel lumen at 50 s post injection and was detected in the ISVs (**Figure 1E**, **Movie 1**). 40 kDa dextran signal in the EVS became more prominent over time, with distal staining at the neuro-muscular junctions after 4.87 min. At 12.1 mpi, 40 kDa dextran signal was detected at the ISV-adjacent myotomal boundaries. This corresponds with eventual detection of 40 kDa dextran in individual skeletal myofiber membranes at approximately 50 mpi (**Figure S1A**). 2000 kDa dextran signal remains in the lumen of the blood vasculature throughout this period, up to 60 mpi in this study (**Figure 1F**). This places fluorophore-labelled 2000 kDa dextran as a suitable tracer control.

**Figure 1.**
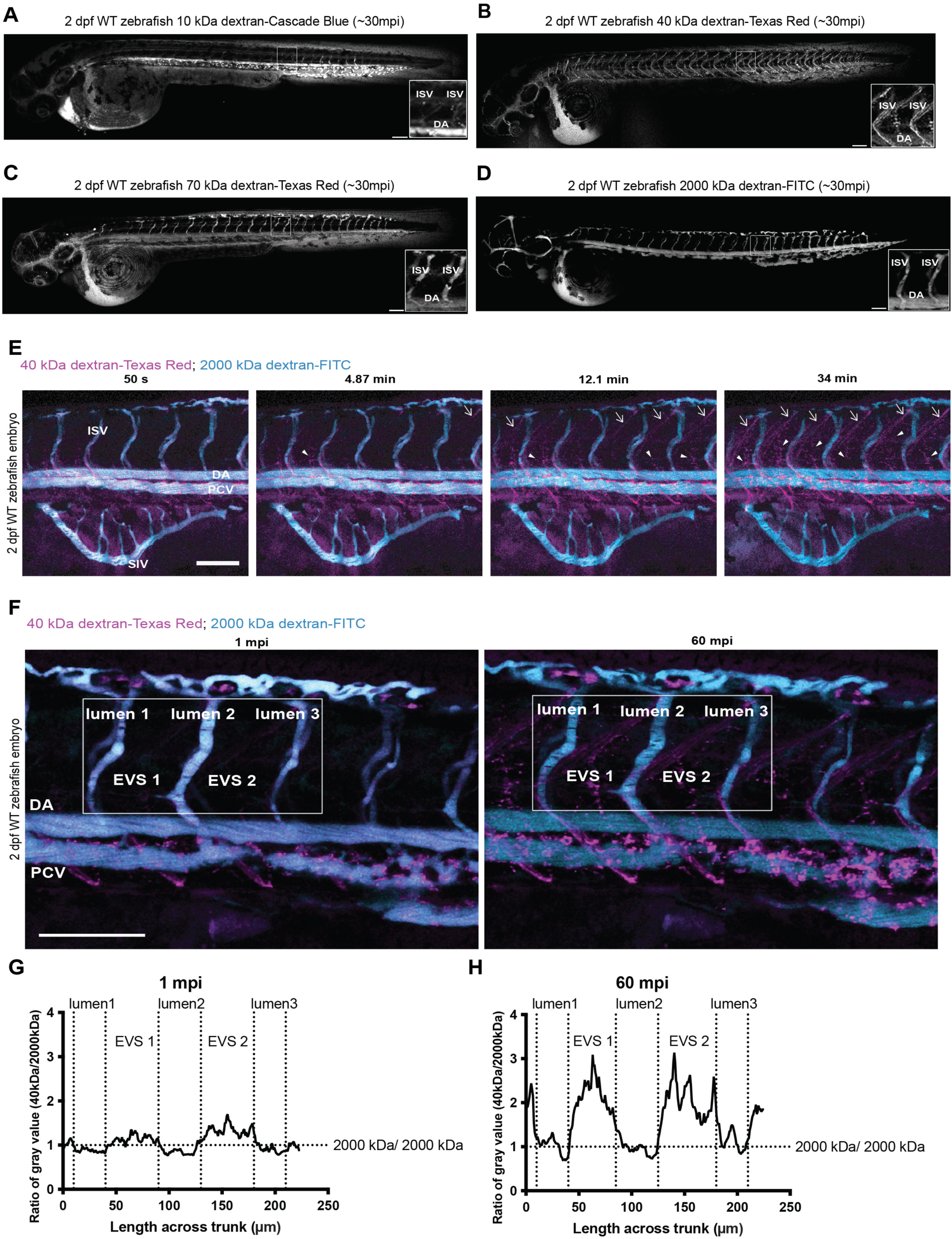
Biodistribution and vascular localization of dextran conjugates microinjected into 2 dpf zebrafish embryos to identify a tracer control and nanoparticle surrogate for studying trans- endothelial transport. **(A-D)** 2 dpf WT zebrafish embryos injected with 10- 2000 kDa dextran conjugates and imaged at approximately 30 mpi. Insets are magnified images of the DA and two ISVs. Z projected images. **(E)** 2 dpf zebrafish injected with 40 kDa dextran-Texas Red as a nanoparticle surrogate and 2000 kDa dextran-FITC as a tracer control and imaged over a period of 50 s to 34 mpi (based on Z projected time series in **Movie 1**). Arrowheads indicate neuromuscular junctions. Arrows indicate myotome boundaries. **(F)** Z projected image of a 2 dpf zebrafish injected with 40 kDa dextran-Texas Red and 2000 kDa dextran-FITC at 1 and 60 min post injection with a consistent region of interest (ROI) (boxed in white) covering two segments of extravascular space (EVS 1-2) in between lumens of intersegmental vessels (lumen 1-3) highlighted. **(G-H)** Column average plots of the ROI in **(F)** showing the signal of 40 kDa dextran-Texas Red corrected with the baseline signal of 2000 kDa dextran-FITC. Scale bars: **A-F**, 100 μm. ISV=intersegmental vessel; EVS=extravascular space; DA=dorsal aorta; PCV=posterior cardinal vein.

To develop a standardized framework for assessing trans-endothelial transport, 40 kDa dextran was employed as a surrogate for a fluid-phase nanoparticle pool. 2 dpf zebrafish embryos were injected with 40 kDa dextran-Texas Red and 2000 kDa dextran-FITC and imaged at 1 and 60 mpi (**Figure 1F**). Quantification was performed using two complementary methods, both normalized to the baseline signal provided by 2000 kDa dextran. First, a column average plot was applied to calculate vertically averaged pixel intensities across two main EVS regions, located between three ISV blood vessel lumens within the same embryo (**Figure 1F**). This standardized region of interest (ROI), defined relative to the yolk extension, was selected due to its inclusion of permeable ISVs and a substantial area of the EVS (**Figure 1F**). ^41, 42^ This analysis revealed increased 40 kDa/2000 kDa signal ratio in EVS 1 and EVS 2 over 60 minutes, indicating extravasation of 40 kDa into the EVS (**Figure 1G-H**). Second, an unbiased quantification method was developed using an ImageJ macro script. This signal measurement script leverages the 2000 kDa dextran signal to threshold and segment the vascular lumen, which also allows for EVS identification via inversion of the thresholded lumen (**Figure 2A-H**). The 2000 kDa dextran signal also serves as a baseline for ratiometric and standardized quantitation of nanoparticle signal within the ISV lumen and surrounding EVS (**Figure 2D-I**). Using this script, quantitative analysis on injected 2 dpf WT embryo images showed a significant increase in the ratiometric signal of 40 kDa dextran-Texas Red to 2000 kDa dextran-FITC in the EVS from 1 to 60 mpi (**Figure 3A**, **Figure S1B**). No significant change in 40 kDa dextran signal was observed within the lumen, likely due to saturation from the injected dextran at this timepoint (**Figure 3A**, **Figure S1B**). These data established the 2000 kDa dextran as a reliable tracer control. This approach reliably controls for injection variability for each microinjection of individual embryos. Collectively, the combined use of 2000 kDa dextran tracer control and quantitation methods, such as the column average plot analysis and the ImageJ signal measurement macro script, provides complementary approaches to reliably detect and quantify the extravasation of 40 kDa dextran from the vascular lumen to the EVS.

**Figure 2.**
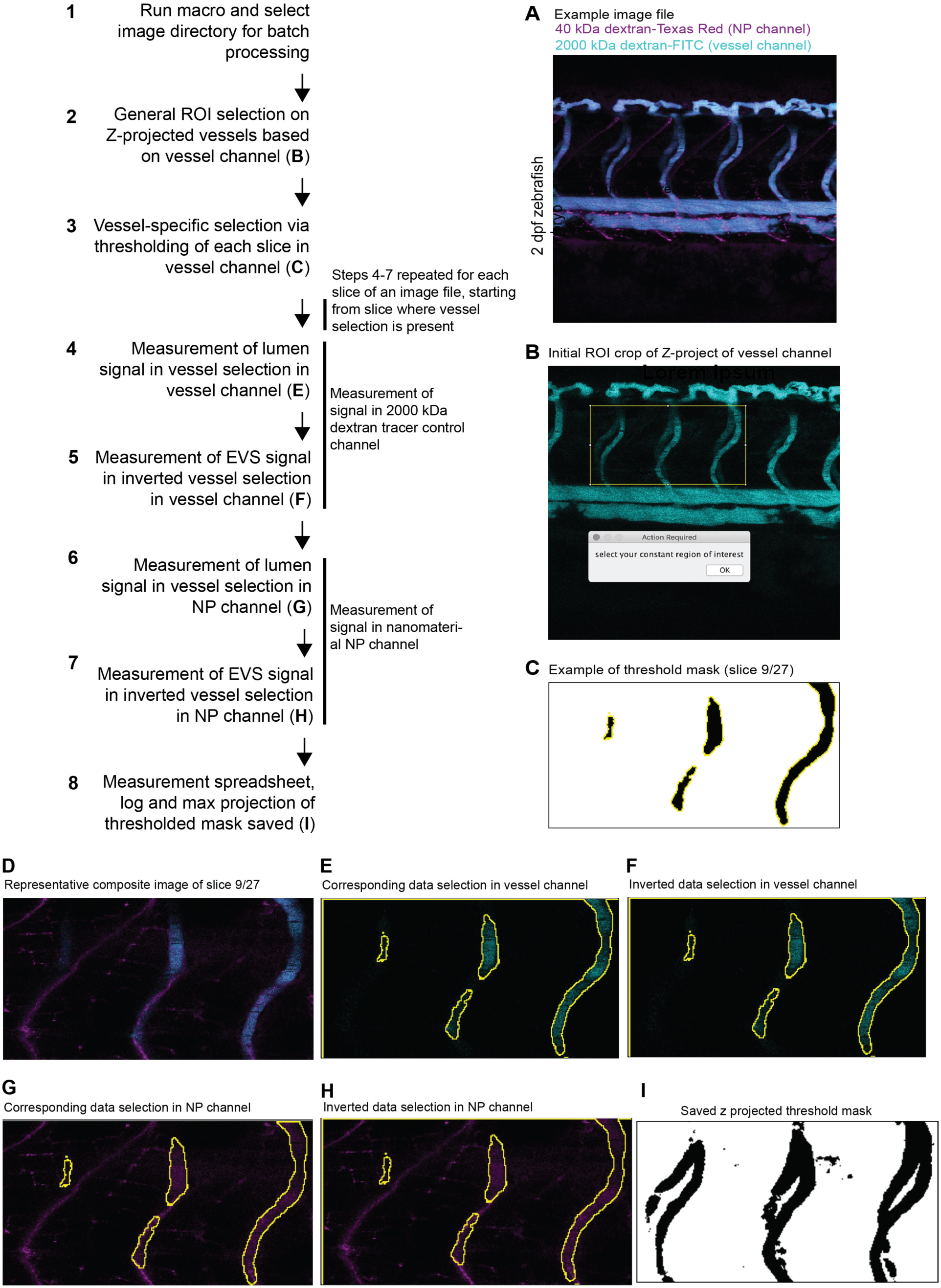
Nanoparticle signal measurement macro script for unbiased quantitation of trans- endothelial transport. When started, the script will prompt the user to select an image directory for batch processing. Selecting an image directory will open single image files containing a ‘vessel channel’ and a ‘Nanoparticle (NP) channel’ which displays signal from circulating 2000 kda dextran and nanoparticle of interest respectively.**(A)** shows an example image file with 40 kDa dextran-Texas Red as a nanoparticle surrogate in the NP channel. Z-projection of the vessel channel **(B)** will be automated to guide the location of embryo and the user will be prompted to manually select a consistent ROI. The script will then perform specific vessel selection by thresholding each Z slice in the vessel channel [example slice in **(C)**]. Starting from the slice where vessel selection is present, for each slice present in the image file, lumen and EVS signal is measured in the vessel selection and inversion of the vessel selection respectively **(D-F)**. The measurements in the vessel channel are the baseline/background values for quantitation provided by the circulating 2000 kDa dextran-FITC. The lumen and EVS signal measurements are then repeated in the NP channel using the same vessel channel selections [represented by **(G-H)**], which provides the nanoparticle signal. The Z-projection of the thresholded vessel is then saved **(I)**, alongside the measurement spreadsheet and log.

**Figure 3.**
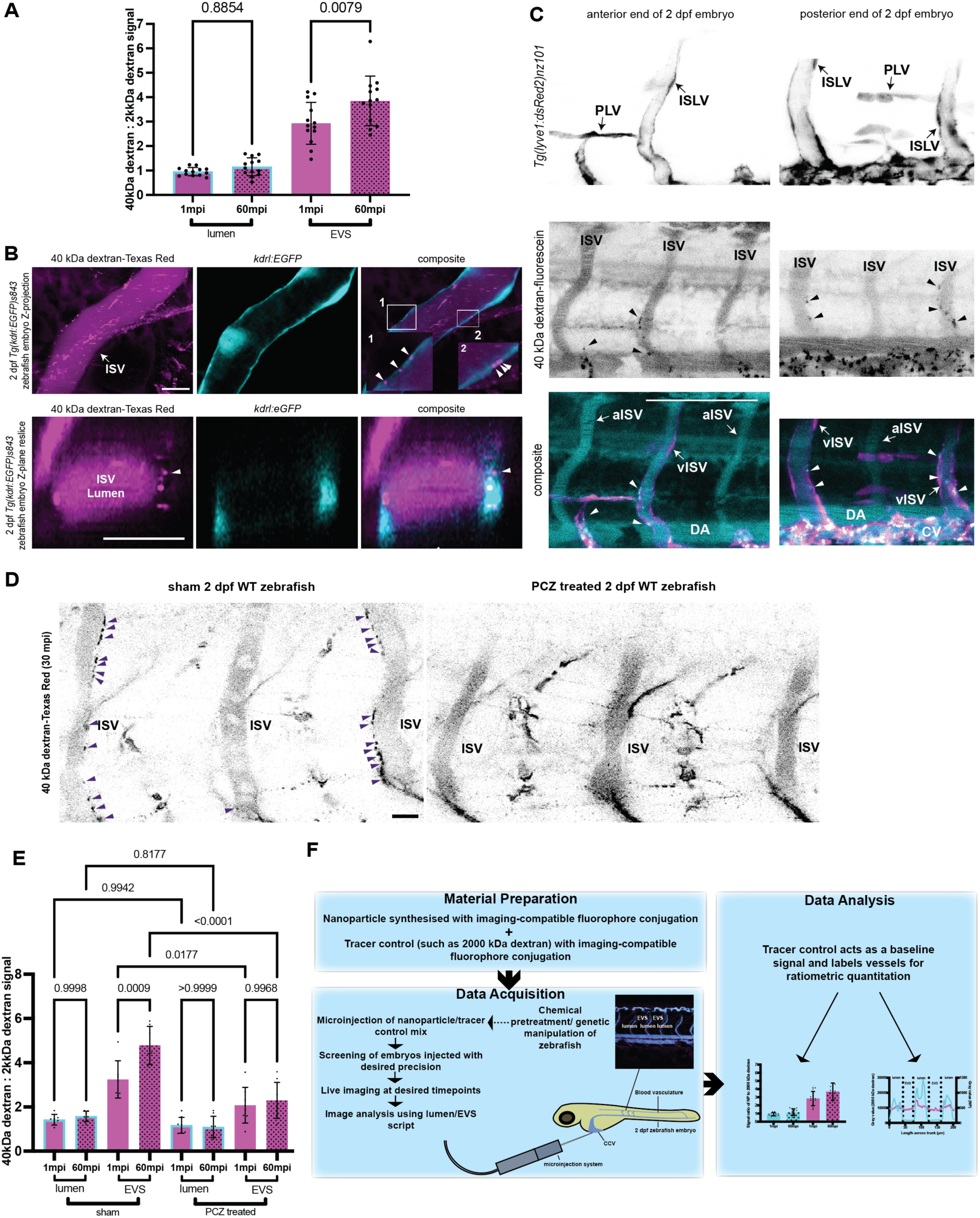
Characterization of the physiological trans-endothelial transport of 40 kDa dextran as a nanoparticle surrogate in the zebrafish via chemical pretreatment. **(A)** Ratiometric signal of 40 kDa dextran-Texas Red (2000 kDa dextran-FITC as baseline) in the lumen and EVS of WT 2 dpf zebrafish at 1 and 60 mpi. Fish number=13; mixed clutch of 6. **(B)** Live images of, and Z-plane reslice through a ISV of 2 dpf *Tg(kdrl:EGFP)^s843^* zebrafish injected with 40 kDa dextran-Texas Red. **(C)** Representative images of the ISVs at the posterior and anterior end of the same 2 dpf *Tg(lyve1:dsRed2)^nz101^* embryo injected with 40 kDa dextran-fluorescein. **(D)** Representative 30 mpi live images of sham and prochlorperazine (PCZ) treated WT 2 dpf zebrafish injected with 40 kDa dextran-Texas Red. Arrowheads indicate dynamic vesicles along the ISVs. **(E)** Ratiometric signal of 40 kDa dextran-Texas Red (2000 kDa-FITC as baseline) in the lumen and EVS of sham treated and PCZ treated WT zebrafish at 1 and 60 mpi. Fish number=7-8; mixed clutch of ≥5 for each group. **(F)** Workflow schematic depicting a standardized protocol to study nanoparticle biodistribution and trans-endothelial trafficking in the zebrafish. A tracer control such as 2000 kDa dextran conjugated to a fluorophore is co-administered with the nanoparticle of interest. This allows for standardization during and after microinjection of the nanoparticle/tracer control mix into embryos pretreated with chemicals and inhibitors. The tracer control allows for screening of embryos injected with desired accuracy and acts as a baseline signal for ratiometric quantitation. ISV=intersegmental vessel; EVS=extravascular space. Quantitation: **A, E**, one-way ANOVA with Tukey’s multiple comparisons test; numeric P values displayed. Data are presented as mean±SD. Scale bars: **B**, **D**, 10 μm. **C**, 100 μm. ISLV=intersegmental lymphatic vessel; PLV=parachordal lymphatic vessel; aISV=arterial intersegmental vessel, vISV=venous intersegmental vessel; ISV=intersegmental vessel; DA=dorsal aorta; CV=caudal vein.

### Uncovering the Trans-endothelial Routes taken by Fluid-phase Dextran in the Early Stages of Circulation via a Standardized Protocol

As a proof of concept for the viability of this standardized protocol, we characterized the physiological trans-endothelial transport pathway taken by the 40 kDa dextran surrogate in the live zebrafish. Live confocal imaging revealed dynamic 40 kDa dextran-positive vesicles at the ISV of injected *Tg(kdrl:EGFP)^s843^* zebrafish embryos (**Figure 3B**, **Movie 2**). Colocalization of 40 kDa dextran with specific GFP expression in endothelial cells of injected *Tg(kdrl:EGFP)^s843^* transgenic zebrafish suggests that these vesicles were intracellular (**Figure 3B**). ^43, 44^ Using *Tg(lyve1:dsRed2)^nz101^* transgenic zebrafish to label venous ISV, ^45, 46^ we showed that these intracellular vesicles were not stochastically distributed across all ISVs of the embryo, but were specifically localized to venous ISVs, and not arterial ISVs (**Figure 3C**).

We hypothesized that these vesicles are endocytic. To test this, we used a pretreatment protocol in which 2 dpf zebrafish were first injected via the CCV with specific dynamin-dependent endocytosis inhibitors prochlorperazine (PCZ) ^47–49^ or dyngo4a. ^2, 50^ After 30 min, pre-treated zebrafish were then injected with the 40 kDa/2000 kDa dextran solution alongside the specific pre-treatment inhibitor and imaged for 60 min. All PCZ-treated embryos possessed a complete loss of dynamic and intracellular vesicles at the ISV (**Figure 3D**). Between 1 mpi and 60 mpi, sham control embryos showed a significant increase in 40 kDa dextran signal at the EVS, indicating extravasation (**Figure 3E**, **Figure S2A**). This extravasation was not observed in PCZ-treated embryos, which possessed no significant difference between 40 kDa dextran signal in the EVS at timepoints 1 mpi and 60 mpi, and significantly lower 40 kDa dextran signals in the EVS compared to untreated embryos (**Figure 3E**, **Figure S2A**). This difference is mirrored in dyngo4a-treated embryos, in which 40 kDa dextran signal in the EVS was significantly reduced and extravasation was lost in dyngo4a-treated embryos (**Figure S2B-C**). Thus, using a combination of live imaging and our signal measurement script, we demonstrated that steady state trans-endothelial trafficking of 40 kDa dextran as a fluid-phase tracer involves dynamic, intracellular, and dynamin-dependent endocytosis at the venous ISV.

Taken together, we have established a standardized protocol capable of evaluating physiological trans-endothelial transport in live zebrafish, employing a 2000 kDa dextran tracer control. This tracer is first used to screen embryos for desired precision in injection and intact tissue morphology, and subsequently serves as a baseline signal for quantitative analysis (**Figure 3F**). An inherent limitation of using the tracer control is that signal in the EVS is not necessarily strictly extravascular, as the 2000 kDa dextran, at timepoints below 60 mpi, primarily localizes to the vascular lumen without labelling endothelial cells. Consequently, intracellular vesicles within endothelial cells contribute to the EVS signal. Nevertheless, this standardized protocol remains adequately sensitive to signal variations at the cellular level, particularly in the context of trans-endothelial transport. Importantly, tracking of the 2000 kDa dextran tracer ensures that the signal of intravenously injected nanoparticles originates specifically from the blood vasculature lumen, enabling confirmation of signal changes in the EVS, particularly during chemical manipulation of embryos.

### 3nm and 7 nm PEG-based Hyperbranched Polymers Traverse the Endothelial Barrier via Paracellular Routes

Having validated the standardized protocol for the sensitive detection and quantification of 40 kDa dextran extravasation, we next investigated the trans-endothelial trafficking routes taken by PEG-based nanoparticles in live zebrafish. Hereafter, unless otherwise stated, all assessment of trans-endothelial transport followed the standardized protocol described in **Figure 3F**. The pretreatment protocol was used for all embryo treatments, and all measurements and analyses were ratiometric, utilizing the 2000 kDa dextran signal as a baseline. As a first example of archetypical ‘soft’ PEG-based nanoparticles, two HBPs of 3 and 7 nm in diameter were synthesized following established methods. ^51^ These polymers are cross-linked polymeric structures which comprise a methoxy-PEG (mPEG) surface chemistry and incorporate a Cyanine 5 (Cy5) dye to enable imaging.

Upon injection into 2 dpf zebrafish embryos, the 3 nm and 7 nm PEG-based HBP-Cy5 exhibited diffuse signals throughout the embryo, indicating rapid systemic distribution (**Figure 4A-B**, **Figure S3B-C**). Given their size, we hypothesized that the trans-endothelial transport route of these PEG-based nanoparticles is paracellular. ^35^ Previous studies have identified zebrafish ISV junctions as sites of vascular permeability. ^41, 42^ Endothelial cell-cell adhesion has been shown to be regulated by the barrier-protective cAMP signalling pathway, wherein both direct and indirect elevation of intracellular cAMP levels have been shown to reduce paracellular permeability. ^52–54^ Thus, to investigate whether paracellular transport contributes to the extravasation of 3 to 7 nm PEG-based HBP, 2 dpf zebrafish embryos were pre-treated with membrane-permeable cAMP. After 30 min, cAMP pre-treated embryos were injected with solutions containing 3 or 7 nm HBP-Cy5, 2000 kDa dextran-FITC and membrane-permeable cAMP and imaged immediately. At 1 mpi, in sham controls, 3 and 7 nm HBP-Cy5 were immediately detected in the EVS (**Figure 4B-C**, **E, Figure S3A**). The column average plot revealed that 3 and 7 nm PEG HBP-Cy5 signals were continuous in the vascular lumens and EVS of sham control embryos (**Figure 4C**, **Figure S3D**), indicative of rapid diffusion throughout the embryo. The cAMP-treated embryos exhibited similar levels of 3 nm or 7 nm HBP-Cy5 in the EVS at 1 mpi (**Figure 4D-E**, **Figure S3A, C-F**). Previous studies have suggested the roles of endothelial cell gap formation, adhesion and spreading in regulating vascular permeability, and that due to this, cAMP elevation performed *in situ* and *in vivo* resulted in an attenuated increase in permeability compared to cultured endothelial monolayers. ^54–56^ This discrepancy may account for the comparable levels of 3 and 7 nm PEG-based HBPs observed in the EVS between the cAMP-treated and sham groups, suggesting incomplete efficiency of the cAMP pretreatment in enhancing paracellular barriers in live zebrafish embryos. However, the column average plot showed that in cAMP-treated embryos, 3 and 7 nm PEG HBP-Cy5 closely followed the trend of the 2000 kDa signal specifically within the vascular lumens, indicating accumulation in circulation (**Figure 4D**, Figure S3D). This was further supported by quantitation, which revealed significantly higher levels of 3 nm and 7 nm HBP in the vascular lumens of cAMP-treated embryos compared to sham controls (Figure 4E, **Figure S3A, E-F**). This indicates enhanced confinement of circulating 3 nm and 7 nm HBP-Cy5 to the bloodstream of cAMP-treated embryos. Dynamin-dependent endocytosis inhibition via PCZ pretreatment of 7 nm HBP-injected embryos did not affect the levels of 7 nm HBP-Cy5 in the lumen and EVS compared to sham controls (Figure 4E, **Figure S3A**). This suggests that the rapid extravasation of 3 nm and 7 nm PEG-based HBP predominantly occurs via passive paracellular pathways rather than intracellular mechanisms.

**Figure 4.**
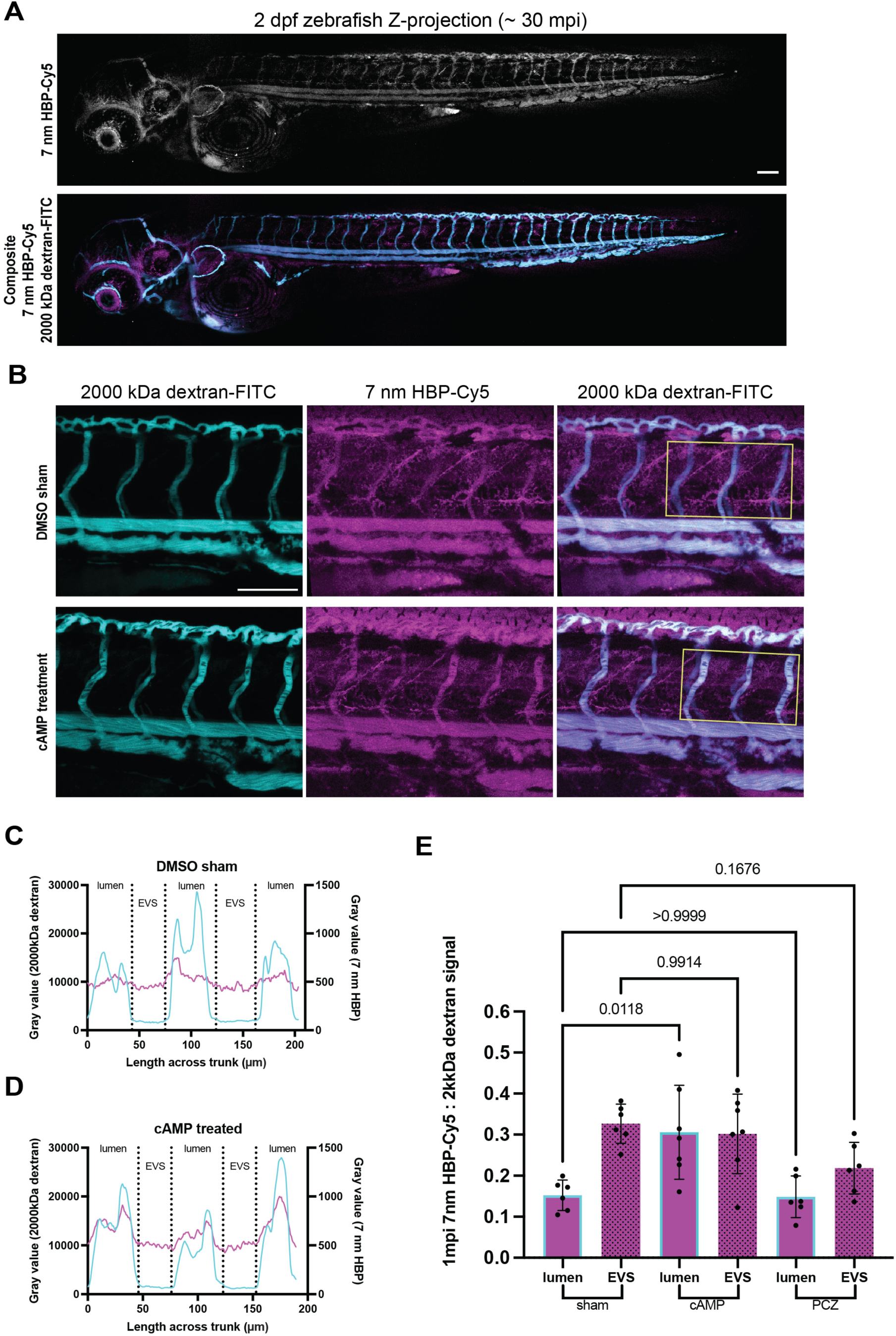
Trans-endothelial transport of 7 nm PEG HBP in live zebrafish. **(A)** Whole mount live images of 2 dpf WT zebrafish injected with 7 nm HBP-Cy5 and 2000 kDa dextran-FITC at approximately 30 mpi. **(B)** Live images of 2 dpf WT zebrafish pre-treated with DMSO or cAMP and injected with 7 nm HBP-Cy5 and 2000 kDa dextran-FITC at 1 mpi. **(C-D)** Column average plots with approximate lumen and EVS boundaries of the ROI in **(B)** showing the signal of 7 nm HBP-Cy5 and 2000 kDa dextran-FITC in 1 WT zebrafish pre-treated with DMSO or cAMP. 7 nm HBP-Cy5 signal in magenta and 2000 kDa dextran-FITC signal in cyan. **(E)** Ratiometric signal of 7 nm HBP-Cy5 (2000 kDa-FITC as baseline) in the lumen and EVS of DMSO (sham), cAMP and PCZ treated WT zebrafish at 1 mpi. Fish number=6-7; mixed clutch of ≥8 for each group. EVS=extravascular space. Quantitation: **E**, one-way ANOVA with Tukey’s multiple comparisons test; numeric P values displayed. Data are presented as mean±SD. Scale bars: **A-B**, 100 μm.

### 32 nm and 47 nm PEG-based Micelles Extravasate from the Blood Vasculature via Dynamin-dependent Endocytosis

To investigate the role of size in the trans-endothelial transport of PEG-based nanoparticles, we then synthesized larger PEG-based micelles of 32 and 47 nm in diameter. ^57^ These micelles were assembled from amphiphilic block-copolymers comprising a hydrophobic vinyl acetate, vinyl bromobutanoate, 2-methylene-1,3-dioxepane copolymer core and a hydrated mPEG hydrophilic corona. After azidation of the bromine-containing monomer, DBCO-functional Cy5 was incorporated into the core of these polymers prior to crosslinking to produce stable polymeric micelles for further study. These micelles can be directly compared to PEG-based HBPs used above due to their equivalent mPEG hydrated surface and spherical shape. Ensuring the surface chemistry of all nanoparticles studied was equivalent (mPEG) enabled us to directly probe the impact of size without varying other physicochemical properties of the nanoparticles.

At 1 mpi, following administration into the bloodstream of 2 dpf zebrafish embryos, the 32 nm PEG micelle-Cy5 was primarily restricted in the blood vessels (Figure 5A). At 60 mpi, 32 nm PEG micelle-Cy5 signal was observed in the EVS, indicating extravasation (Figure 5A). We hypothesized that these 32 nm micelles were trafficked in endocytic compartments, as dynamic and intracellular vesicles were observed in the ISV endothelial cells (**Movie 3**). Zebrafish embryos pre-treated with PCZ were injected with 32 nm PEG micelle-Cy5. At 60 mpi, PCZ-treated embryos possessed a significantly higher signal in the lumen and significantly lower signal in the EVS compared to sham controls (Figure 5B, **Figure S4A**). Dynamic vesicles carrying 32 nm PEG micelle-Cy5 signal in the ISV endothelial cells were also reduced (**Movie 4**). Strikingly, cAMP-treated zebrafish embryos injected with 32 nm PEG micelle-Cy5 exhibited no significant changes in EVS and lumen signals, compared to sham controls, indicating that the trans-endothelial transport of 32 nm PEG micelle-Cy5 is not dependent on paracellular routes (Figure 5B, **Figure S4A**).

**Figure 5.**
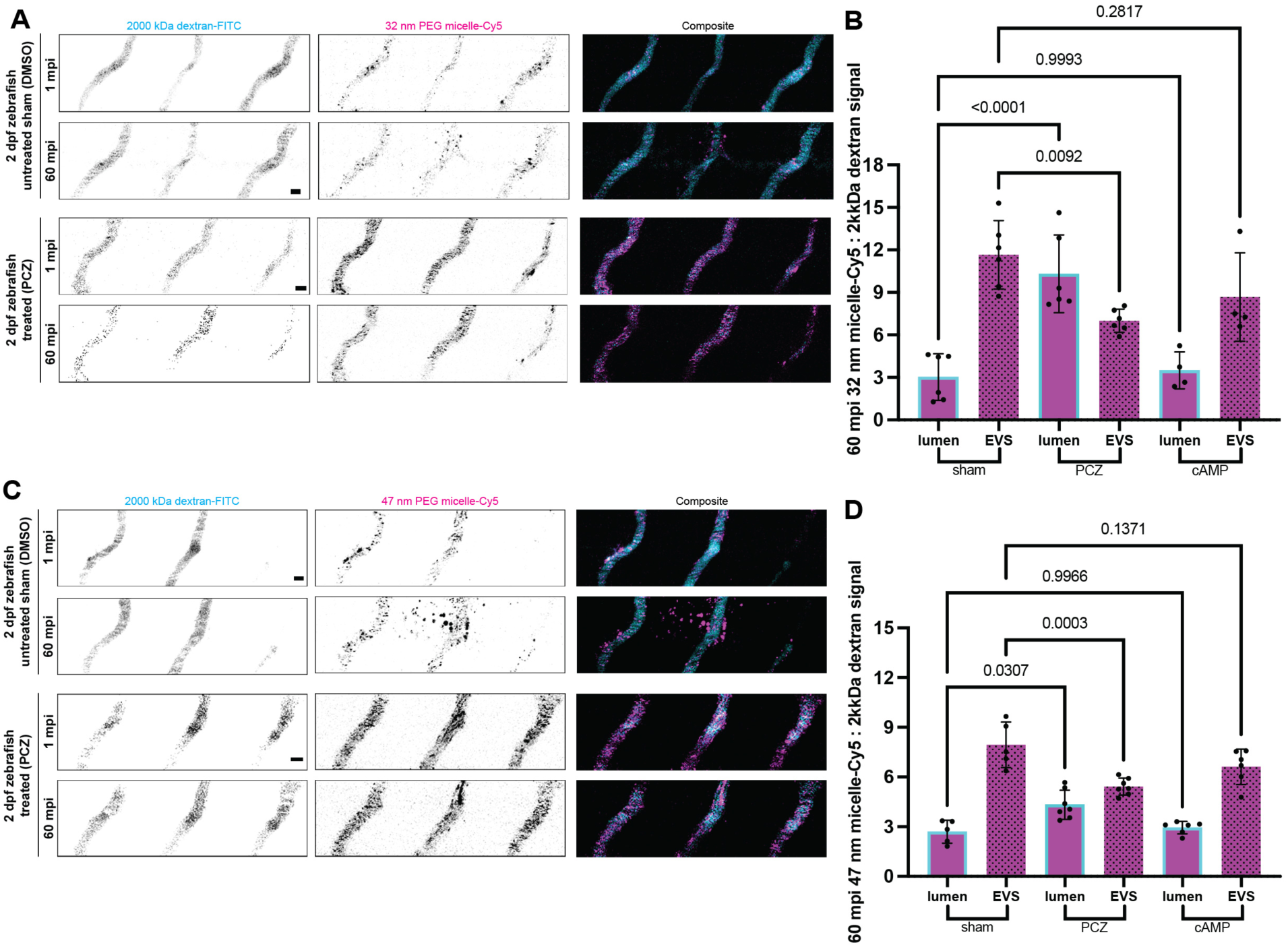
Characterization of the trans-endothelial transport of 32 nm and 47 nm PEG micelles in the zebrafish. **(A)** 2 dpf WT zebrafish pre-treated with DMSO (sham control) or PCZ, and injected with 32 nm PEG micelle-Cy5 and 2000 kDa dextran-FITC at 1 and 60 mpi. **(B)** Ratiometric signal of 32 nm PEG micelle-Cy5 (2000 kDa-FITC as baseline) in the lumen and EVS of DMSO (sham), PCZ and cAMP treated WT zebrafish at 60 mpi. Fish number=4-6; mixed clutch of ≥8 for each group. **(C)** 2 dpf WT zebrafish pre-treated with DMSO (sham control) or PCZ and injected with 47 nm PEG micelle-Cy5 and 2000 kDa dextran-FITC at 1 and 60 mpi. **(D)** Ratiometric signal of 47 nm PEG micelle-Cy5 (2000 kDa-FITC as baseline) in the lumen and EVS of DMSO (sham), PCZ and cAMP treated WT zebrafish at 60 mpi. Fish number=5-7; mixed clutch of ≥6 for each group. Data are presented as mean±SD. Quantitation: **B**, **D** one-way ANOVA with Tukey’s multiple comparisons test; numeric P values displayed. Scale bars: **A**, **C** 10 μm.

Administration of the larger 47 nm PEG micelle-Cy5 into 2 dpf zebrafish embryos resulted in a biodistribution pattern comparable to that of 32 nm PEG-micelle-Cy5, with signal initially confined to the blood vessels and extravasation to the EVS at 60 mpi (**Figure 5C**, **Figure S4B**). 47 nm PEG micelle-Cy5 injected embryos pretreated with PCZ exhibited confinement of the nanoparticles in blood vessels (**Figure 5C**). Quantitation on 47 nm PEG micelle-Cy5 injected embryos revealed that PCZ-treated embryos possessed a significantly higher signal in the lumen and lower signal in the EVS compared to sham controls, while cAMP-treated embryos possessed no significant differences in levels of 47 nm PEG micelle-Cy5 in the lumen and EVS (**Figure 5D**, **Figure S4B**). Taken together, these findings demonstrate that following intravenous administration, PEG-based micelles of 32 nm to 47 nm in diameter extravasate via sequestration into dynamic vesicles associated with dynamin-dependent endocytosis. This extravasation does not involve inter-endothelial, paracellular transport. This sheds light on the relatively uncharacterized mechanisms underlying steady-state trans-endothelial transport and transcytosis of nanoparticles. ^58, 59^ In epithelial models, the dynamin-dependent, clathrin-mediated pathway is involved in transcytosis, however, its role in endothelial cells remains poorly explored. ^58–61^ Previous work has demonstrated and discussed the entry of sub-50 nm particles such as ferritin into clathrin-coated pits in endothelial cells of 5 dpf zebrafish via bulk phase endocytosis. ^13^ Notably, this parallels the characteristics of specific tumor endothelial cells (termed nanoparticle transport endothelial cells, N-TECs) which mediate nanoparticle transport into solid tumors. ^29^ Transcriptomic analysis revealed a high upregulation of the clathrin-mediated pathway in N- TECs, suggesting that clathrin-mediated transport may serve as a mechanistically relevant route for trans-endothelial transport. It is important to note that other endocytic pathways cannot be ruled out, particularly the FEME pathway, which has been shown to be inhibited by PCZ. ^47, 62^ Altogether, these data suggest that, to a significant extent, bulk phase endocytosis characterized by dynamin-dependent endocytic vesicles constitutes a primary mechanism underlying the extravasation of 32-47 nm PEG-based nanoparticles in the live zebrafish.

### Trans-endothelial Transport of PEG-based Nanoparticles is Unaffected by Caveolae In Zebrafish and Mouse Models

Finally, we used the standardized protocol to test the contribution of caveolae to trans- endothelial transport *in vivo*. The transport of nanoparticles across the endothelium is not well-characterized, but caveola and its components are often implicated. ^13, 32, 58, 59^ Inhibitors of caveola internalization have been shown to be widely unspecific, making genetic knockout of caveola components the preferred approach to study caveola roles, particularly in a live organism. ^2, 13^ The formation of caveolae requires the expression of various caveola-associated proteins such as Caveolin1 and Cavin1 which associates primarily with the caveola bulb. ^2, 31^ The zebrafish expresses Cavin1a and Cavin1b as Cavin1 paralogs. ^63, 64^ We employed two previously generated zebrafish lines - a knock-in zebrafish line expressing endogenous Cavin1b-mScarlet (*TgKI(cavin1b-mScarlet)^pd126^*) and a complete Cavin1 double knockout (DKO) zebrafish line (*cavin1a/1b* DKO line). ^65, 66^ We first characterized Cavin1b expression in the developing zebrafish by imaging the *TgKI(cavin1b-mScarlet)^pd126^* line. At 2 dpf, Cavin1b-mScarlet was detected in the ISV, dorsal aorta (DA) and dorsal longitudinal anastomosing vessel (DLAV) (**Figure 6A**, **Figure S5A**). This expression pattern persists throughout the 8 dpf stage, at which point a complex vasculature network begins to form (**Figure S5A**). ^45, 67^ At 15 dpf, Cavin1b-mScarlet was detected throughout most of the vasculature, including superficial intersegmental blood vessels (SIV), intercostal vessels (ICV) and the caudal fin vascular network (**Figure 6B**, **Figure S5A**). Interestingly, under confocal microscopy, Cavin1b-mScarlet expression was detected in the DA but not in the posterior cardinal vein (PCV) across developmental stages up to 15 dpf (**Figure 6B**, **Figure S5A**). This is possibly due to differences in flow and tension accommodation between these two vessel types. ^68, 69^ Caveola structures have been previously observed in the endothelial cells of mice and zebrafish embryos. ^13, 70^ Several early studies implicate a role for caveolae in mediating transcytosis of macromolecules and endothelial permeability. ^71–73^ However, there are discrepancies in caveola-deficient models, in which normal or heightened permeability of caveola-null blood vessels have been described. ^74, 75^ This was attributed to the concurrent opening of paracellular pathways, in which Cavin1 and Caveolin1 KO mice exhibited discontinuous endothelial cells separated by gaps, and a reduction in the size of endothelial cell-cell tight junctions respectively. ^70, 75, 76^ Transmission electron microscopy (TEM) identified abundant caveolae in the blood vessel of 17 dpf WT juveniles, comparable to the abundance of caveolae in the gastrocnemius muscle capillary of adult WT mice (**Figure S5B- C**). Genetic knockout of Cavin1a and Cavin1b, and Cavin1 in zebrafish and mouse models respectively resulted in a marked reduction in caveola density within these vessels (**Figure S5B- C**). Notably, no defect in endothelial cell-cell junctions was observed in 17 dpf *cavin1a/1b* DKO zebrafish juveniles (**Figure S5B**). This places the *cavin1a/1b* DKO zebrafish line as a suitable model to dissect the role of caveolae in trans-endothelial transport without the involvement of paracellular defects.

**Figure 6.**
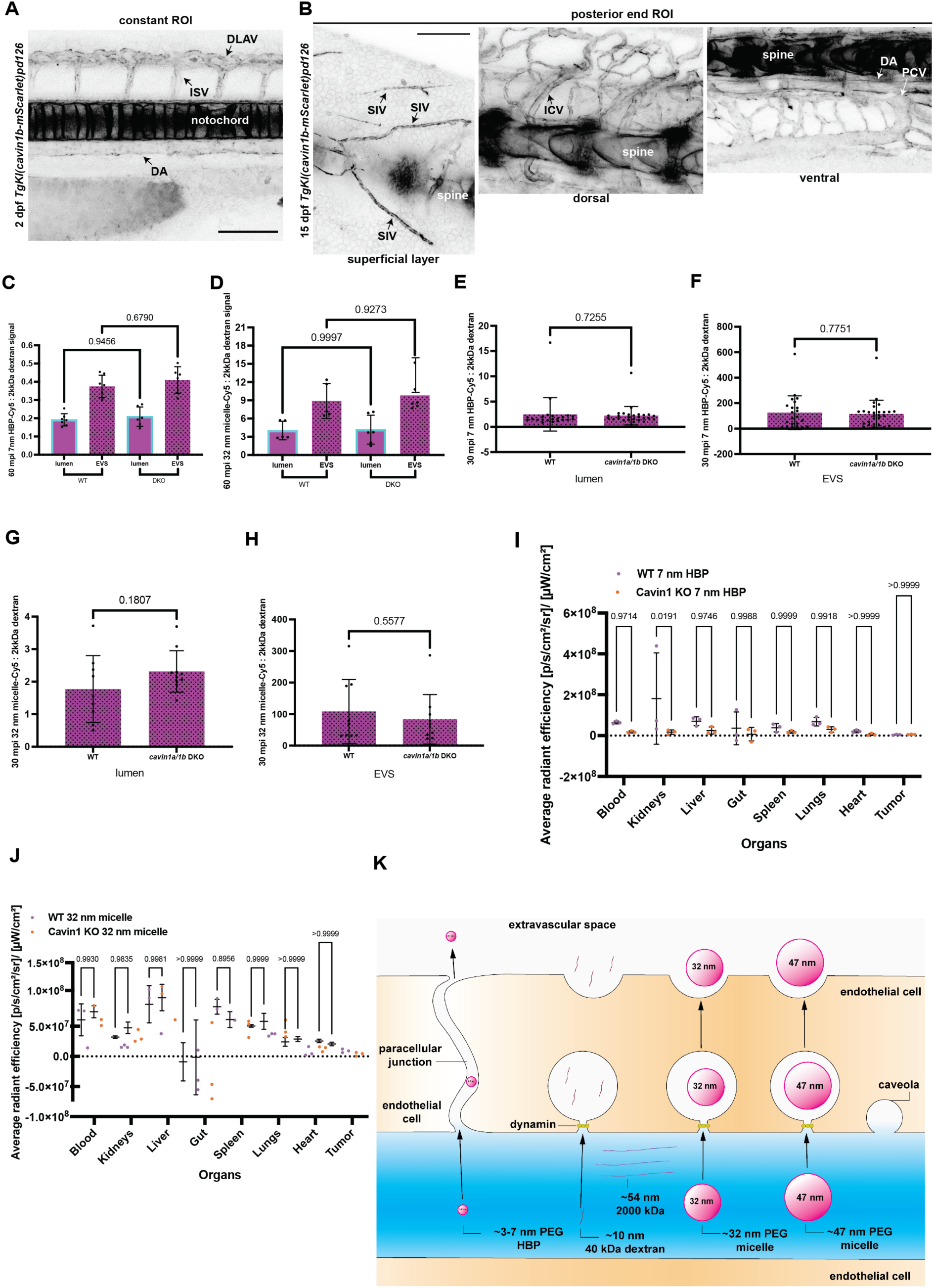
Contribution of caveolae to PEG-based nanoparticle trans-endothelial transport. (**A**) 2 dpf *TgKI(cavin1b-mScarlet)^pd126^* transgenic zebrafish expressing Cavin1b-mScarlet. (**B**) Z projected images of 15 dpf *TgKI(cavin1b-mScarlet)^pd126^* transgenic zebrafish expressing Cavin1b-mScarlet. (**C)** Ratiometric signal of 7 nm PEG HBP-Cy5 (2000 kDa-FITC as baseline) in the lumen and EVS of 2 dpf WT and *cavin1a/1b^-/-^* DKO zebrafish at 60 mpi. Fish number=6- 7; mixed clutch of ≥5 for each group. (**D)** Ratiometric signal of 32 nm PEG micelle-Cy5 (2000 kDa-FITC as baseline) in the lumen and EVS of 2 dpf WT and *cavin1a/1b^-/-^* DKO zebrafish at 60 mpi. Fish number=5-6; mixed clutch of ≥4 for each group. **(E-F)** Ratiometric signal of 7 nm PEG HBP-Cy5 (2000 kDa-FITC as baseline) in the lumen and EVS of 15 dpf WT and *cavin1a/1b^-/-^* DKO zebrafish at 30 mpi. Fish number= 22-27; mixed clutch of ≥7 for each group. **(G-H)** Ratiometric signal of 32 nm PEG micelle-Cy5 (2000 kDa-FITC as baseline) in the lumen and EVS of 15 dpf WT and *cavin1a/1b^-/-^* DKO zebrafish at 30 mpi. Fish number=9-10; mixed clutch of ≥5 for each group. **(I-J)** Quantification of average radiant efficiency in the ROIs of tissues in Cavin1 and WT mice allografted with B16 tumors and administered with Cy5 labeled 7 nm PEG HBP and 32 nm PEG micelles. N=3 for each group. **(K)** Schematics of the trans- endothelial transport routes taken by PEG-based nanoparticles of sizes 3 to 47 nm within the first hour of circulation. PEG-based nanoparticles below 7 nm cross the endothelial barrier via the paracellular junction. As a PEG-based nanoparticle increases in size beyond 7 nm (up to at least 47 nm), active dynamin-mediated endocytic uptake is adopted over paracellular uptake during trans-endothelial transport. Quantitation: **C-D** one-way ANOVA with Tukey’s multiple comparisons test; **E-H** two tailed unpaired t-test; **I-J** two-way ANOVA with Sidak’s multiple comparisons test. Numeric P values displayed. Data are presented as mean±SD. Scale bars: **A-B**, 100 μm. ISV=intersegmental vessel; DA=dorsal aorta; DLAV=dorsal anastomosing vessel; SIV=superficial intersegmental blood vessel; ICV=intercostal vessel; PCV=posterior cardinal vein.

Nanoparticles with PEG modifications have been variably reported to engage with caveolae in their uptake and transcellular transport. ^2, 28, 32, 77–79^ To investigate the role of caveolae in the trans-endothelial transport of our PEG-based nanoparticles, we injected 2 dpf *cavin1a/1b* DKO zebrafish with 7 nm PEG HBP-Cy5 and 32 nm PEG micelle-Cy5. At 60 mpi, Cavin1a/1b DKO zebrafish embryos possessed no significant differences in their levels of 7 nm PEG HBP-Cy5 or 32 nm PEG micelle-Cy5 in the endothelial lumen and EVS compared to WT controls (**Figure 6C-D**, **Figure S6A-B**). This indicates that in 2 dpf embryos expressing relatively low levels of Cavin1 paralogs (**Figure 6A**), trans-endothelial transport of 7 nm PEG HBP-Cy5 and 32 nm PEG micelle-Cy5 does not require caveolae and their protein components. As juvenile zebrafish possessed a high density of caveolae in their endothelial cells (**Figure S5B**), we next injected these PEG-based nanoparticles into 15 dpf zebrafish. The signal measurement script in the standardized protocol effectively parsed the complex vascular network of the juvenile zebrafish using the 2000 kDa dextran tracer control (**Figure S7A-B**). Both 7 nm PEG HBP-Cy5 and 32 nm PEG micelle-Cy5 injected 15 dpf cavin1a/1b DKO zebrafish exhibited no significant difference in their nanoparticle signals in the lumen and EVS compared to WT controls (**Figure 6E-H**). Collectively, these results indicate that caveolae do not contribute significantly to the physiological trans-endothelial transport of PEG-based nanoparticles 7 nm PEG HBP-Cy5 and 32 nm PEG micelle-Cy5 in the zebrafish. This negligible effect of caveola loss on trans- endothelial transport was also reflected in tumor-bearing Cavin1 KO mice administered with these nanoparticles, in which no significant differences in biodistribution were detected (**Figure 6I-J**).

The internalization of nanoparticles via caveolae or association with caveolar proteins remains a disputed topic. ^2, 32^ Previous work has demonstrated that the caveolar lumen in zebrafish endothelial cells can accommodate 5 nm ferritin, suggesting potential nanoparticle packaging into recycling endosomes for transcellular transport. ^13^ However, in the case of the 7 nm PEG HBP-Cy5, rapid extravasation appears to be primarily mediated by paracellular pathways rather than active transport such as bulk-phase endocytosis. By comparison, extravasation of multiple 32 nm PEG micelles would likely necessitate structural or functional modifications to fit the spatial constraints of the sub-50 nm caveolar vesicles, if these nanoparticles were to facilitate their transcellular transport via caveola-associated pathways.

Cumulatively, this study, using the standardized protocol, elucidated the trans-endothelial transport mechanisms of PEG-based nanoparticles ranging from 3 nm to 47 nm within the first hour of intravenous administration (**Figure 6K**). The trans-endothelial transport routes of these PEG-based nanoparticles are dependent on their size. Smaller 3 to 7 nm PEG HBP traverse the endothelial barrier via passive paracellular routes. While uptake by endocytic vesicles may occur, the extravasation of these nanoparticles is disproportionately facilitated by inter-endothelial gaps. As PEG-based nanoparticle size increases to 32 nm and 47 nm, the contribution of paracellular pathways markedly decreases, while bulk-phase endocytosis via dynamin-dependent pathways facilitates transcellular transport. Although caveolar vesicles may accommodate these nanoparticles, caveolae-mediated pathways do not contribute to their extravasation.

## Conclusion

We have established a standardized protocol for studying trans-endothelial transport in live zebrafish. The co-administration of 2000 kDa dextran is a central component of the workflow. We show that this standardized protocol is capable of thresholding complex zebrafish vascular networks, applicable to developmental stages up to 15 dpf. By incorporating a tracer control, this standardized protocol expands upon previous sophisticated methods developed for quantifying nanoparticle biodistribution and circulation in the zebrafish. ^7, 8, 15^ We emphasize the co-administration of a large tracer, such as 2000 kDa dextran, as essential for studies involving nanoparticle administration in zebrafish that require standardized controls. This tracer remains confined within the blood vasculature lumen for a defined period, providing a stable reference for the concentration of the co-injected particle during that timeframe, allowing standardization in individual zebrafish embryos. We have recently investigated blood-brain barrier permeability in the zebrafish following the principle of using a large tracer control, such as 2000 kDa dextran, demonstrating its fundamental applicability. ^80^

Following the workflow of this standardized protocol, we characterized physiological transcellular transport in live zebrafish embryos and revealed the size dependent trans- endothelial transport of PEG-based HBPs and micelles of 3 to 47 nm in diameter. Using knockout and knock-in zebrafish lines alongside mouse models, we demonstrated that caveolae are not required for the extravasation of these PEG-based nanoparticles. The zebrafish lines used in this study are viable, with their blood vasculature representing a model for an endothelial barrier in steady-state conditions. Such models are advantageous for testing various nanoparticles designed for non-leaky vasculature and lymphatics. This study provides fundamental data on nano-bio interactions between PEG-based nanoparticles and zebrafish endothelial cells *in vivo*, enabling further research into biodistribution, half-time and long-term effects of diverse nanoparticles, and targeted delivery in xenografted lines.

## Methods

### 32 nm and 47 nm PEG Micelle-Cy5 Synthesis

The synthesis of amphiphilic micelles has been described previously. ^57^ Below is a summary of synthetic detail, along with characterisation data.

### General procedure for the synthesis of mPEG-*b*-[poly(VAc-*co*-VBr-*co*-MDO)]

MDO (0.126 g, 1.10 × 10^−3^ mol), VAc (0.50 g, 2.60 × 10^−3^ mol), VBr (0.50 g, 2.60 × 10^−3^ mol), 3 (9.20 mg, 3.70 × 10^−5^ mol), ABCN (0.610 mg, 3.70 × 10^−6^ mol), and benzene (66 wt%) were placed into a Young’s tapped ampoule and degassed via freeze-pump-thaw (3x cycles) then backfilled with argon. The resulting solution was stirred and heated to 90 °C for 48 h before the polymerisation was quenched in an ice bath. An aliquot was taken before precipitation to determine the monomer conversion using ^1^H NMR spectroscopy. The polymer was dissolved in a small amount of CHCl^3^ and precipitated twice in ice-cold Et^2^O. The pale-yellow solid was then dried *in vacuo* in a vacuum desiccator over P^2^O^5^ for 24 h.

P1 – [VAc]:[VBr]:[MDO] = 0.4:0.4:0.2; target DP = 250

*M*^n,NMR^ = 24.7 kDa; *M*^n,SEC^ = 15.6 kDa; *Ð*^M^ = 1.56

P2 – [VAc]:[VBr]:[MDO] = 0:0.8:0.2; target DP 250

*M*^n,NMR^ = 27.0 kDa; *M*^n,SEC^ = 14.8 kDa; *Ð*^M^ = 1.61

### General procedure for azidation to mPEG-*b*-[poly(VAc-*co*-VN^3^-*co*-MDO)]

mPEG-*b*-[poly(VAc-*co*-VBr-*co*-MDO)] (0.5 g) was dissolved in DMF (2.5 mL) and stirred for 10 min at RT. To this solution was added sodium azide (10 equivalents per bromide), and the resultant heterogeneous solution was stirred at RT for a further 24 h, upon which the solution turned a dark blue colour. After this time, the solution was poured into CH^2^Cl^2^ (50 mL) and extracted with brine (3 x 50 mL) where the colour proceeded to disappear. The organic phase was then collected, dried over anhydrous MgSO^4^, filtered and reduced in volume. The polymer was then precipitated from a minimum amount of CHCl^3^ into ice-cold diethyl ether, centrifuged (5 min @ 5000 RCF, 4 °C) to separate the solid from the supernatant, and then the solid was dried *in vacuo* in a vacuum desiccator over P^2^O^5^ for 24 h to yield a very pale-yellow solid. SEC (THF) *M*^n^ = 10.0 kDa, *Ð*^M^ = 1.20; ATR-FTIR (ν/cm^−1^) 2095 (N^3^ azide stretch).

### Preparation of SPAAC crosslinked micelles

mPEG-*b*-[poly(VAc-*co*-VN^3^-*co*-MDO)] (10 mg) was dissolved in 2 mL of MeCN in a glass vial. To this, Cy5-DBCO was added (0.1 molar equivalent per chain), and the solution was stirred at RT overnight. To this solution, 18 ml of Milli-Q water was added slowly at a rate of 1.8 ml/h via a syringe pump injector, and the solution was stirred with vigorous agitation. After addition, DBCO-PEG^4^-DBCO (0.5 equivalents to azide in MeCN) was quickly added to this solution and stirred for a further 24 h. After this time, the solution was dialysed against Milli-Q water in 3.5 kDa MWCO Snakeskin dialysis tubing at room temperature for 3 days with frequent water changes. The solution was then filtered through a 0.45-micron nylon filter ready for use.

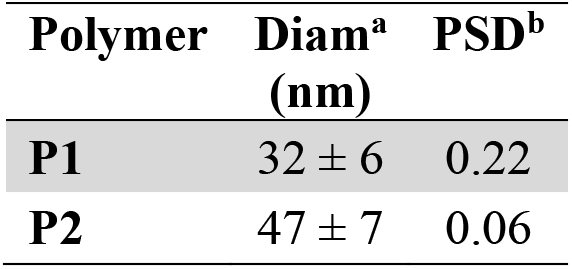

a % number distribution; b Particle Size Distribution.

### 3 nm and 7 nm PEG HBP-Cy5 Synthesis

3 nm and 7 nm PEG HBP-Cy5 were produced following previously reported methods to produce 3 and 7 nm diameter materials. ^51^

## Animal Models

### Zebrafish Lines

All zebrafish experiments were approved by the University of Queensland (UQ) Animal Ethics Committee: Project Number: 2022/AE000666 and the UQ Biosafety Committee. Zebrafish were raised and maintained according to institutional guidelines (14-h light/10-h dark cycle, The University of Queensland, UQ). Zebrafish embryos up to 5 dpf were raised in standard E3 media (5mM NaCl, 0.17mM KCl, 0.33 mM CaCl^2^, 0.33 mM MgSO^4^) in 8 cm Petri dishes; late-larval to juvenile zebrafish (6 dpf to 45 dpf) were housed in 1 L tanks with flow at 28.5 °C; adult zebrafish (90 dpf above) were housed in 3 or 8 L tanks with flow at 28.5 °C. ^81^ All post-embryonic zebrafish measurements were carried out between tanks of similar population densities and conditions. The following zebrafish strains were used in this study: wild type (WT) TAB, an AB/TU line derived from the Zebrafish International Resource Centre Oregon; ^82^ *cavin1a/1b DKO*, a previously generated double knockout line generated by crossing *cavin1a^-/- uq10rp^* zebrafish to cavin1b^-/-uq7rp^ zebrafish and incrossing the offspring to homozygosity; ^65^ *TgKI(cavin1b-mScarlet)^pd126^*; ^66^ *Tg(lyve1:dsRed2)^nz101^*; ^45^ *Tg(kdrl:EGFP)^s843^*. ^44^ The developmental stages of zebrafish used in experiments are specifically stated in the corresponding figure legends. All reagents were obtained from Sigma-Aldrich unless otherwise specified.

### Zebrafish Intravenous Microinjection and Live Imaging

#### Live imaging of 2 dpf and 15 dpf zebrafish

Zebrafish embryos or juveniles were anesthetized in ethyl 3-aminobenzoate methanesulfonate (tricaine) solution ^81^ until inactivity was determined via touch (after approximately 1 minute). Live 2 to 15 dpf zebrafish were then mounted in 0.8 % low gelling temperature (LGT) agarose solution on MatTek glass bottom dishes in a lateral view unless otherwise stated (anterior to the left, posterior to the right) and submerged in tricaine solution. Mounted live zebrafish were then imaged at 28 °C under a Zeiss LSM880 inverted confocal microscope, using Plan-Apochromat 20x/0.8 NA dry or 40x/1.3 NA oil immersion objectives. Image acquisition was performed using Zeiss ZEN 2012 (black edition) software.

#### 2dpf zebrafish embryos

For intravenous microinjection of 2 dpf zebrafish embryos, embryos were firstly anesthetized in tricaine solution until inactivity was determined (∼1 min), mounted in 0.8 % LGT agarose solution in tricaine on MatTek glass bottom dishes in a lateral view unless otherwise stated (anterior to the left, posterior to the right) and submerged in tricaine solution. ^65^ Microinjection needles (OD 1.0 mm, ID 0.78 mm, length 150 mm, Harvard Apparatus) were pulled with a Sutter P-97 micropipette puller and loaded with the appropriate solution (as detailed below) and calibrated for 1 nL injection volume. Calibration was achieved via injection into mineral oil under an Olympus SZX7 stereo microscope and calculation of volume via the radius of the bolus, as guided by a stage micrometer and eyepiece graticule. Mounted live 2 dpf zebrafish embryos were then microinjected via the common cardinal vein (CCV) and screened for intravenous injection accuracy via 2000 kDa dextran-FITC imaging under the EVOS FL inverted fluorescence microscope (AMG). Screened embryos were then imaged under a Zeiss LSM880 confocal microscope as described above.

#### 15 dpf zebrafish embryos

For intravenous microinjection of 15 dpf juvenile zebrafish, juveniles were first anesthetized in tricaine solution until inactivity was determined (∼1 min) and transferred to a MatTek glass bottom dish. The tricaine solution was then removed, and 2 % LGT agarose solution was immediately applied on the juveniles. The juvenile zebrafish were then re-positioned in a lateral view (anterior to the left, posterior to the right) and incubated at room temperature (RT) for 1 min for solidification of the 2 % LG agarose. Mounted juveniles were then submerged in tricaine solution, and agarose material around the head of the juveniles was removed via forceps. Microinjection needles were pulled, loaded with the appropriate injection solution as detailed below and calibrated (as described above) for an injection volume of approximately 30 nL. Cannulation was achieved via injection into the dorsal aorta above the gills of the juvenile zebrafish. 15 dpf juvenile zebrafish were microinjected with 60 nL of the appropriate solution. Next, microinjected juveniles were screened for intravenous injection accuracy via 2000 kDa dextran-FITC imaging under the EVOS FL inverted fluorescence microscope (AMG). At 30 min post injection (mpi), screened juveniles were then imaged under a Zeiss LSM880 confocal microscope as described above.

#### Nanoparticle and dextran microinjection solution preparation

For all experiments involving the intravenous microinjection of zebrafish embryos and juveniles, all zebrafish were injected with a solution containing the following individual materials with these specific final concentrations: 40 kDa dextran-Texas Red (final concentration of 1.5 mg/mL, Thermo Fisher Scientific) or 40 kDa dextran-fluorescein (final concentration of 1.5 mg/mL, Thermo Fisher Scientific) or 10 kDa dextran-Cascade Blue (final concentration of 1.5 mg/mL, Thermo Fisher Scientific) or 70 kDa dextran-Texas Red (final concentration of 1.5 mg/mL, Thermo Fisher Scientific) or 3 nm PEG HBP-Cy5 (final concentration of 3 mg/mL) or 7 nm PEG HBP-Cy5 (final concentration of 3 mg/mL) or 32 nm PEG micelle-Cy5 (final concentration of 2.368 mg/mL) or 47 nm PEG micelle-Cy5 (final concentration of 2.816 mg/mL). Unless otherwise stated, all solutions used contained 2000 kDa dextran-FITC as a tracer control (final concentration of 1.5 mg/mL), phenol red for brightfield visibility (10% of solution) and E3 media as a diluent.

#### Zebrafish pretreatment protocol

For 2 dpf zebrafish embryos undergoing the pretreatment protocol, embryos were prepared for microinjection as above. Embryos were anesthetized in tricaine, mounted in 0.8 % LGT agarose and intravenously microinjected via the CCV with 1 nL of a solution containing 10 % phenol red, and prochlorperazine (PCZ, final concentration of 6 mM diluted in fresh DMSO) or dyngo4a (final concentration of 10 mM diluted in fresh DMSO, purchased from Selleck Chemicals, Houston, TX, USA) or pCPT-cAMP (final concentration of 15 mg/mL diluted in E3 media). Sham control groups for PCZ, dyngo4a and cAMP treated zebrafish were injected with 1 nL of 40 % DMSO in E3 media, 40 **%** DMSO in E3 media and E3 media respectively. After 30 min, the pretreated embryos were then injected again via the CCV with 1 nL of a solution containing the individual material of interest (final concentration described above), 10 % phenol red and PCZ, dyngo4a or cAMP solution as a diluent with a final concentration of 1.5 mM, 5 mM and 2.5 mg/mL respectively. Sham control groups for PCZ, dyngo4a and cAMP treated zebrafish were injected with 1 nL of a solution containing the individual material of interest (final concentration described above), 10 % phenol red, and E3 or DMSO diluent depending on corresponding treatment group.

#### Mouse *Ex Vivo* Biodistribution Experiment

All animal experiments were approved by the UQ Animal Ethics Committee: Project Number: 2021/AE000860 and UQ Biosafety Committee. *Cavin1*-/- mice, generated by replacing part of Exon 1 of the *cavin1* gene with a LacZ/Neo fusion cassette, were originally obtained from Boston University School of Medicine. ^83^ C57BL/6J [STR9] control mice were acquired from the Animal Resource Centre and were allowed access to food and water *ad libitum* throughout the course of the experiment. Four to seven-month-old *Cavin1*-/- mice and age-matched WT controls (C57BL/6J [STR9]) were inoculated with 250,000 B16 cells and tumors were allowed to develop for approximately 21 days, until tumors were 200 to 500 mm^3^ in size. Cavin1-/- and WT mice were then injected intravenously via the tail vein with 7 nm PEG HBP-Cy5 and 32 nm PEG micelle-Cy5 at concentrations of 10-15 mg/kg for micelles and 25 mg/kg for HBP. Mice were culled 24 hours post administration of the nanoparticles, and their organs were imaged *ex vivo* via an IVIS Lumina X5 imaging system. The average radiant efficiency of Cy5 signals of each organ was calculated via the IVIS software and statistical analysis was performed using Graphpad Prism 10.

#### Electron Microscopy

Electron microscopy of 17 dpf juvenile *zebrafish* and mouse gasteronecmius muscle were performed as described previously. ^84^ Briefly samples were fixed in 2.5% glutaraldehyde then underwent a series of post fixation and contrasting stains in a BioWave microwave (PELCO) at 80W for 3min with vacuum for each step. These were 1.5% potassium ferricyanide and 2% osmium tetroxide in ddH^2^O, 1% thiocarbohydrazide in ddH^2^O, 2% osmium tetroxide in ddH^2^O, lead aspartate (20mM lead nitrate, 30mM aspartic acid, pH 5.5), and 1% uranyl acetate in ddH^2^O with 3 ddH^2^O washes in between each step. Samples then underwent serial dehydration in increasing concentrations of ethanol before infiltration with resin and polymerisation. Ultrathin sections were attained on a UC6 ultramicrotome (Leica) and imaged on a transmission electron microscope (Jeol-1011) at 80kV.

### Imaging and Quantitation

#### Lumen and EVS Nanoparticle Signal Measurement Macro Script

Image processing and quantitation were performed using a custom-developed Fiji macro script (macro available at https://doi.org/10.5281/zenodo.15308422). ^85^ The workflow of the macro is summarized in Figure 2A-I. Upon running the script, the macro prompts the user to open a directory of images to be analyzed. This macro was designed to process image files with the .czi extension, comprising standardized images of microinjected zebrafish (oriented anterior to the left, posterior to the right; with end of yolk extension visible), with the first channel imaged for vascular 2000 ka dextran-FITC (referred to as the ‘green’ or ‘vessel’ channel) and second channel imaged for the material of interest (referred to as the ‘red’ or ‘NP’ channel). A maximum intensity projection of the full Z-stack of each image was then generated to aid in the manual selection of the constant region of interest (ROI) using the rectangle select tool, subsequently cropping the original Z-stack. The constant ROI was defined as a region containing 3 intersegmental vessels (ISV)s, with one ISV located above the tip of the yolk extension. For vessel segmentation, a median filter was applied to the vessel (green) channel and a global threshold was manually set by the user (typically with the lower threshold > 1), generating a binary mask. This mask was then applied slice-by-slice across the Z-stack. For each optical slice, the vessel region (defined by the mask) was measured for mean fluorescence intensity and area in both the vessel (green) and NP (red) channels. The extravascular space (EVS) was defined by inverting the mask, and the mean fluorescence intensity and area were similarly measured. If no vessel region was detected in a slice, that particular slice is excluded from measurements. All output files, including summary files .txt and .xls formats with per-slice measurements of mean fluorescence intensity and area for the green and red channels within vessel and EVS regions, binary masks for each stack, and the corresponding maximum intensity projection of the mask were saved into a new timestamped ‘Results’ subfolder generated within the selected working directory.

For quantification of the injected material relative to the baseline provided by the 2000 kDa dextran-FITC signal, ratiometric analyses were performed on each slice of the image Z-stack. Using outputs generated by the macro above, the integrated densities of the material of interest (red channel) and vessel baseline (green channel, 2000 kDa dextran-FITC) were calculated independently for both the vessel and extravascular space (EVS) regions. Integrated density was defined as the product of mean fluorescence intensity and area for the particular ROI selected within each slice. Ratios of integrated density between the red and green channels were then computed separately for the vessel (lumen) and EVS regions across each slice in the Z-stack. For each image (representing one zebrafish), the mean red-to-green integrated density ratio across all slices was calculated for the vessel and EVS separately, generating a single ‘lumen’ value and a single ‘EVS’ value respectively per fish. Statistical comparisons were conducted using GraphPad Prism 10 (GraphPad Software). Ordinary one-way analysis of variance (ANOVA) with Tukey’s multiple comparisons test was applied on these datasets for 2 dpf experiments. For 15 dpf datasets, unpaired two-tailed t-tests were used separately for lumen and EVS values.

#### Zebrafish study imaging analyses

For column average plots, the Plot Profile function in Fiji was used on rectangular selections of the ROI described above (region containing three ISVs, with one ISV located above the tip of the yolk extension) on images of microinjected zebrafish. Profile plots of fluorescence intensity across the ROI were generated and the resulting data were exported and plotted using GraphPad Prism 10.

## Author Contributions

Conceptualization, Y.-W.L. and R.G.P.; Methodology, Y.-W.L., C.A.B., N.L.F., N.D.C., D.Y.A., J.R., C.F., N.M., J.H., H.P.L., Y.W., Z.H.H., and T.E.H.; Formal Analysis, Y.-W.L.; Writing - Original Draft, Y.-W.L.; Writing - Review & Editing, Y.-W.L., T.E.H., A.K.L., K.J.T. and R.G.P.; Supervision, K.J.T. and R.G.P.; Funding Acquisition, R.G.P., K.J.T.

## Acknowledgments

We thank the members of the Thurecht and Parton groups for helpful discussions, and Michel Bagnat for providing us with the *TgKI(cavin1b-mScarlet)^pd126^* zebrafish line. R.G.P. was supported by an Australian Research Council (ARC) Laureate Fellowship (FL210100107).

N.D.C. is supported as a CZI Imaging Scientist by grant number 2020-225648 from the Chan Zuckerberg Initiative DAF, an advised fund of Silicon Valley Community Foundation. Confocal microscopy was performed at the Australian Cancer Research Foundation (ACRF)/Institute for Molecular Bioscience (IMB) Dynamic Imaging Facility for Cancer Biology, established with funding from the ACRF. The authors acknowledge the facilities, and the scientific and technical assistance, of the Australian Microscopy & Microanalysis Research Facility at the Centre for Microscopy and Microanalysis, The University of Queensland. The authors declare no conflicts of interest.

## Supporting Information

The following files are available free of charge - the Supporting Information includes additional experimental data and details, including Z-projected confocal images of each microinjected zebrafish analyzed by the signal measurement macro, with corresponding threshold masks generated by the signal measurement macro.

**Movie 1**: Time series showing Z projected images of 2 dpf zebrafish injected with 40 kDa dextran-Texas Red and 2000 kDa dextran-FITC and imaged over a period of 50 s to 34 mpi.

**Movie 2**: Signal of circulating 40 kDa dextran-Texas Red in an ISV.

**Movie 3**: ISVs of a sham control zebrafish embryo injected with 32 nm PEG micelle-Cy5.

**Movie 4**: ISVs of a PCZ-treated zebrafish embryo injected with 32 nm PEG micelle-Cy5.

