## Supporting Information for "A Standardized Protocol to Investigate Trans- Endothelial Trafficking in Zebrafish: Nano-bio Interactions of PEG-based Nanoparticles in Live Vasculature"

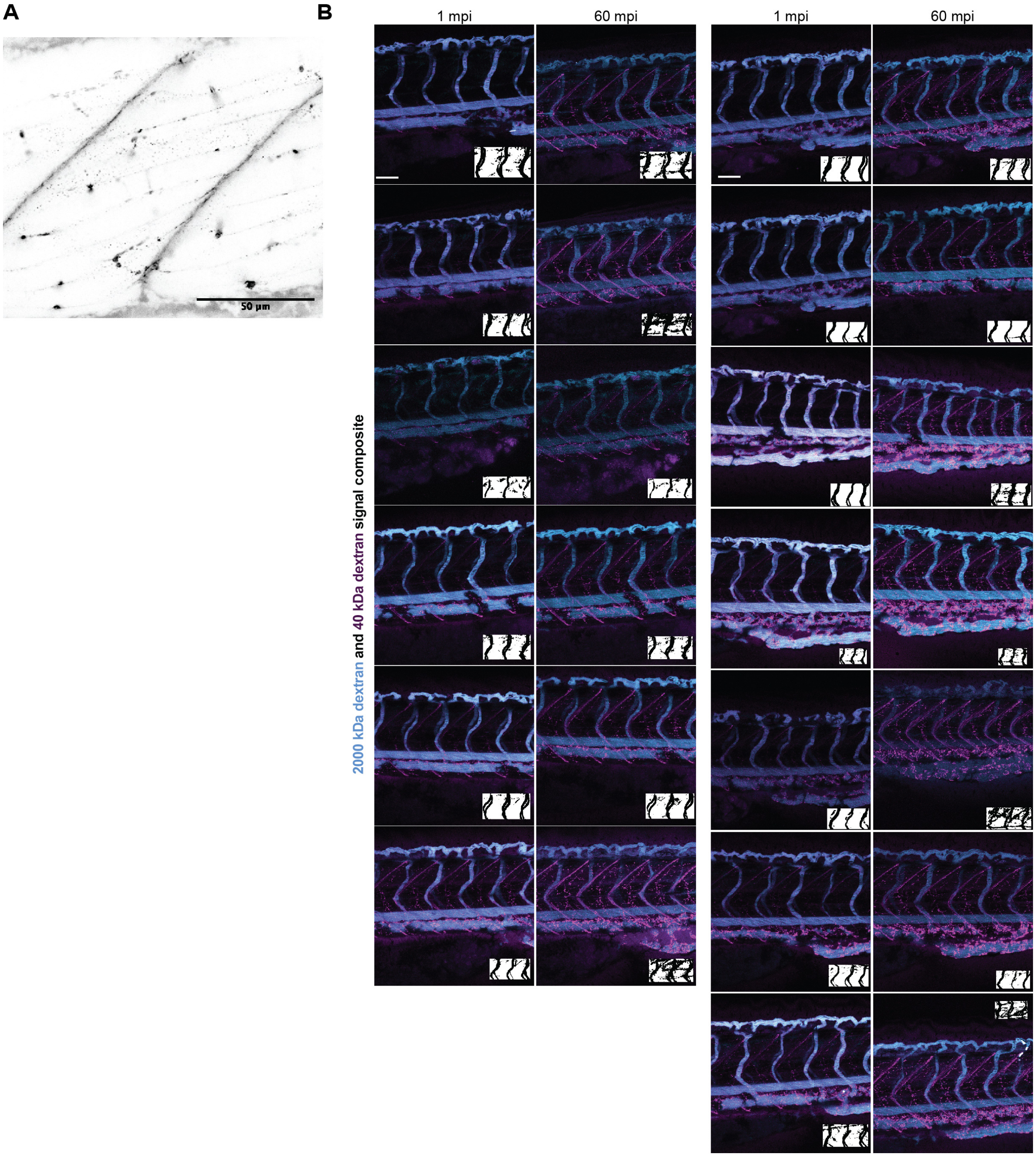


**Figure S1**. (**A)** Skeletal myofiber confocal image of 2 dpf WT zebrafish injected with 40 kDa dextran-Texas Red as a nanoparticle surrogate. Scale bar: 50 µm (**B)** Z-projected confocal images of each microinjected 2 dpf zebrafish analyzed in **Figure 3A**, with insets showing corresponding threshold masks generated by the signal measurement macro. Scale bar: 100 µm

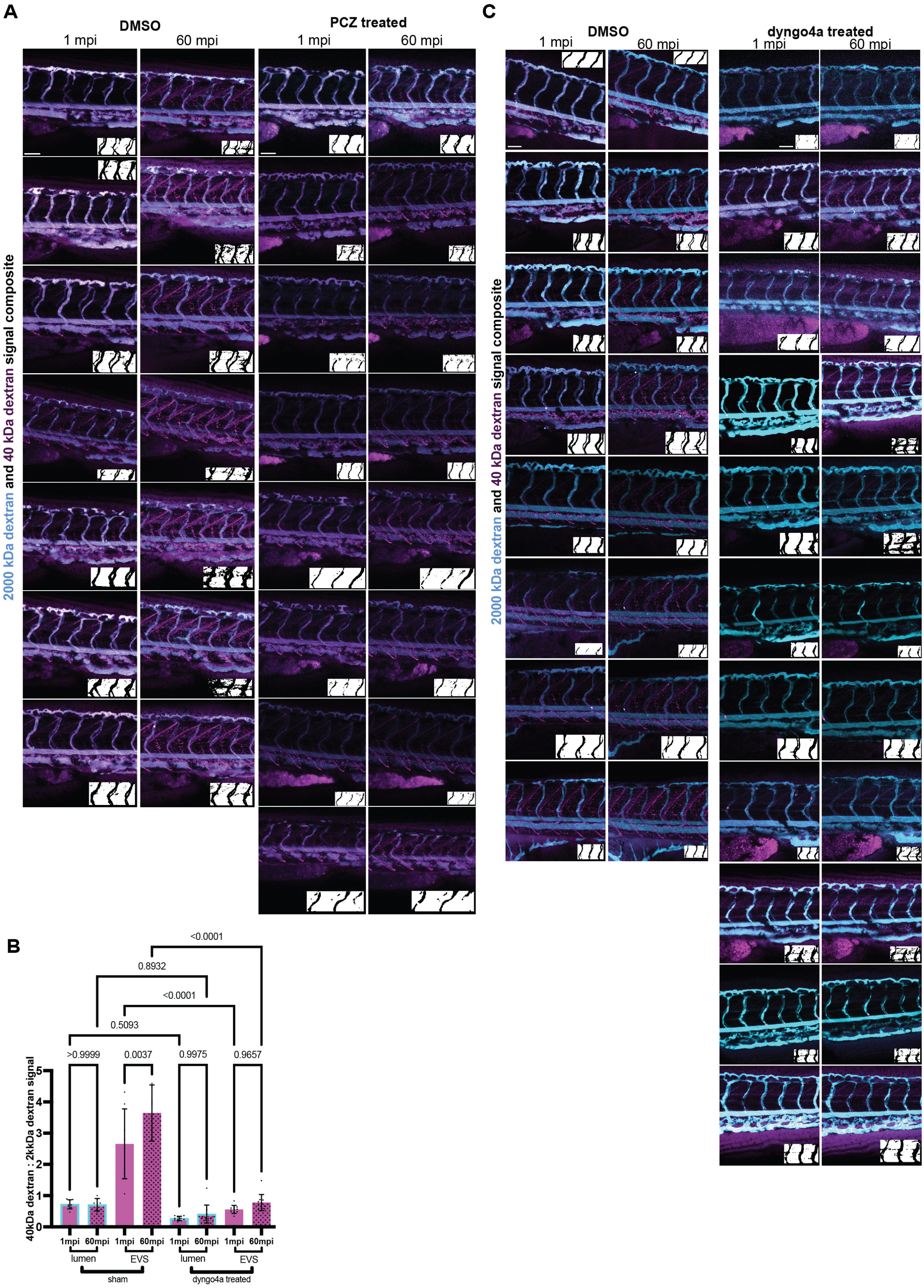


**Figure S2. (A)** Z-projected confocal images of each microinjected 2 dpf zebrafish analyzed in **Figure 3E**, with insets showing corresponding threshold masks generated by the signal measurement macro. Scale bar: 100 µm (**B)** Ratiometric signal of 40 kDa dextran-Texas Red (2000 kDa-FITC as baseline) in the lumen and EVS of sham treated and dyngo4a treated WT zebrafish at 1 and 60 mpi. Fish number 8-11; mixed clutch of ≥10 for each group. Quantitation: one-way ANOVA with Tukey's multiple comparisons test; numeric P values displayed. Data are presented as mean±SD. **(C)** Z-projected confocal images of each microinjected 2 dpf zebrafish analyzed in **(B)**, with insets showing corresponding threshold masks generated by the signal measurement macro. Scale bar: 100 µm

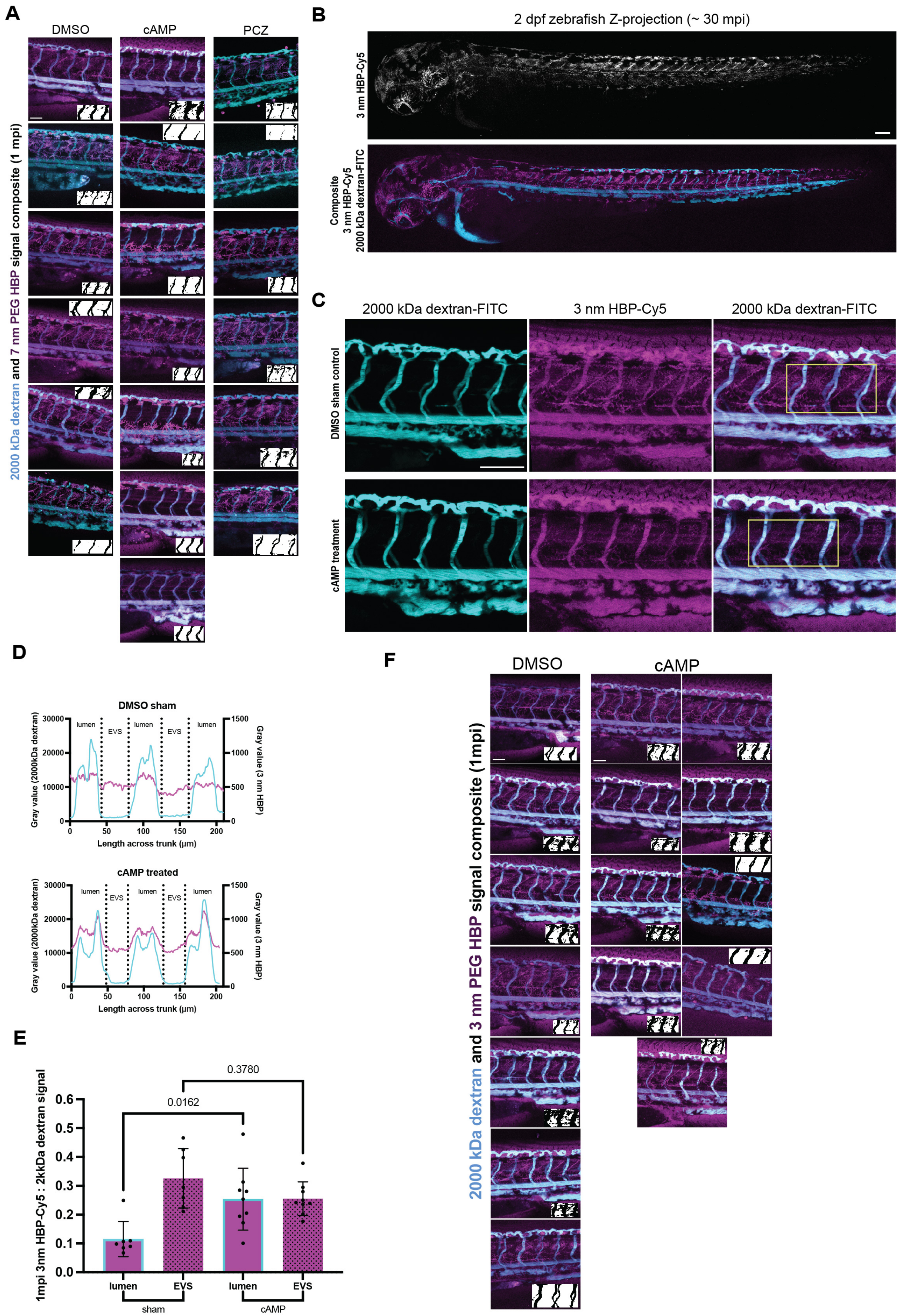

A
DMSO
cAMP
PCZ
2000 kDa dextran and 7 nm PEG HBP signal composite (1 mpi)
B
2 dpf zebrafish Z-projection (~ 30 mpi)
3 nm HBP-Cy5
Composite
3 nm HBP-Cy5
2000 kDa dextran-FITC
C
2000 kDa dextran-FITC
3 nm HBP-Cy5
2000 kDa dextran-FITC
DMSO sham control
cAMP treatment
D
DMSO sham
cAMP treated
lumen
EVS
Gray value (2000kDa dextran)
Gray value (3 nm HBP)
Length across trunk (µm)
E
1mpi 3nm HBP-Cy5 : 2kkDa dextran signal
0.0162
0.3780
lumen
EVS
sham
cAMP
F
DMSO
cAMP
2000 kDa dextran and 3 nm PEG HBP signal composite (1mpi)

**Figure S3. (A)** Z-projected confocal images of each microinjected 2 dpf zebrafish analyzed in **Figure 4E**, with insets showing corresponding threshold masks generated by the signal measurement macro. Scale bar: 100 µm **(B)** Whole mount live images of 2 dpf WT zebrafish injected with 3 nm HBP-Cy5 and 2000 kDa dextran-FITC at approximately 30 mpi. Scale bar: 100 µm **(C)** Live images of WT zebrafish pre-treated with DMSO (sham control) or cAMP and injected with 3 nm HBP-Cy5 and 2000 kDa dextran-FITC at 1 mpi. Yellow boxes indicate constant ROI. Scale bar: 100 µm **(D)** Column average plots with approximate lumen and EVS boundaries of the ROI in **(C)** showing the signal of 3 nm HBP-Cy5 and 2000 kDa dextran-FITC in 1 WT zebrafish pre-treated with DMSO or cAMP. **(E)** Ratiometric signal of 3 nm HBP-Cy5 (2000 kDa-FITC as baseline) in the lumen and EVS of DMSO sham treated and cAMP treated WT zebrafish at 1 mpi. Fish number=7-9; mixed clutch of ≥8 for each group. **(F)** Z-projected confocal images of each microinjected 2 dpf zebrafish analyzed in **(E)**, with insets showing corresponding threshold masks generated by the signal measurement macro. Scale bar: 100 µm

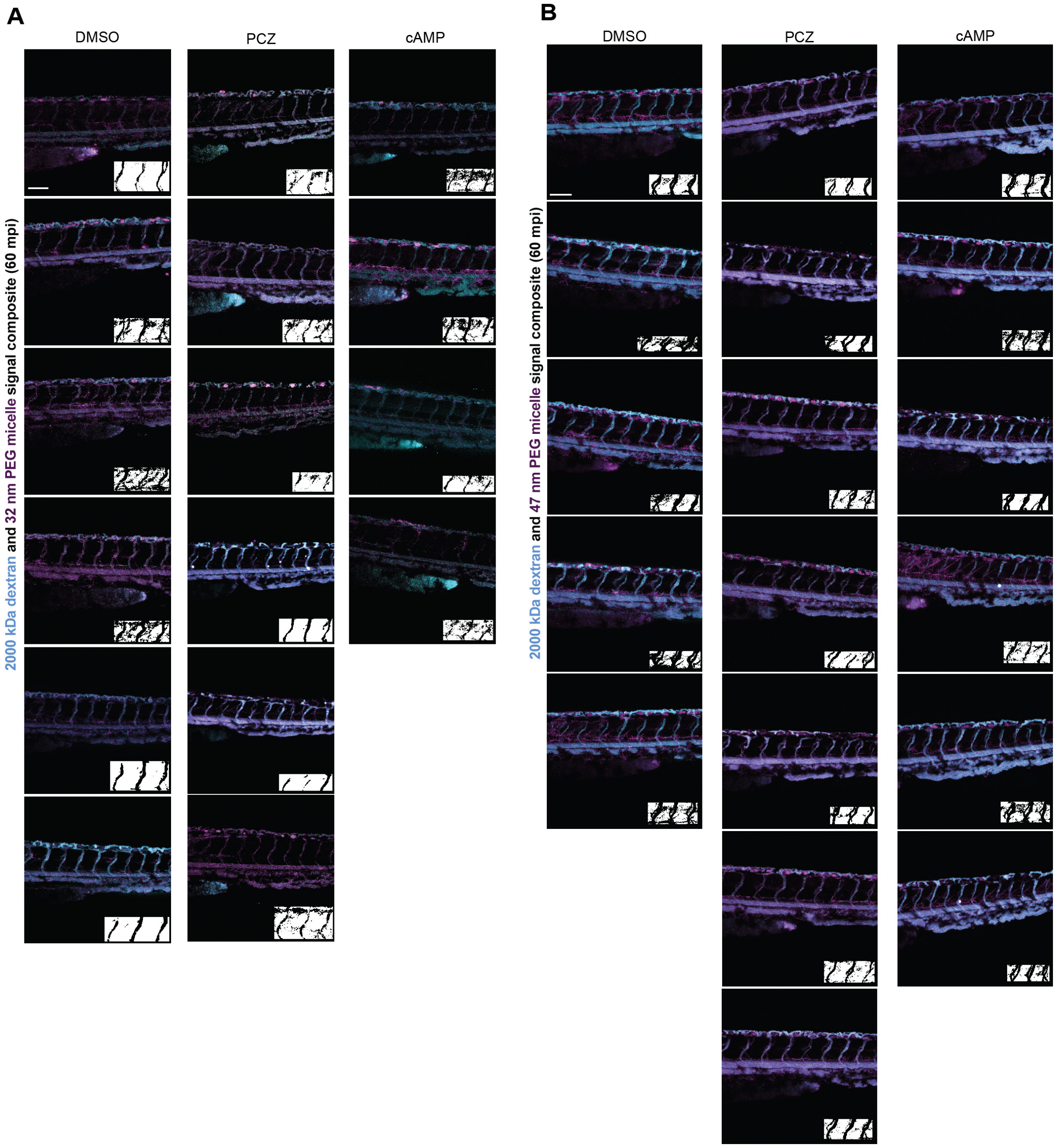


**Figure S4. (A)** Z-projected confocal images of each microinjected 2 dpf zebrafish analyzed in **Figure 5B**, with insets showing corresponding threshold masks generated by the signal measurement macro. Scale bar: 100 µm **(B)** Z-projected confocal images of each microinjected 2 dpf zebrafish analyzed in **Figure 5D**, with insets showing corresponding threshold masks generated by the signal measurement macro. Scale bar: 100 µm

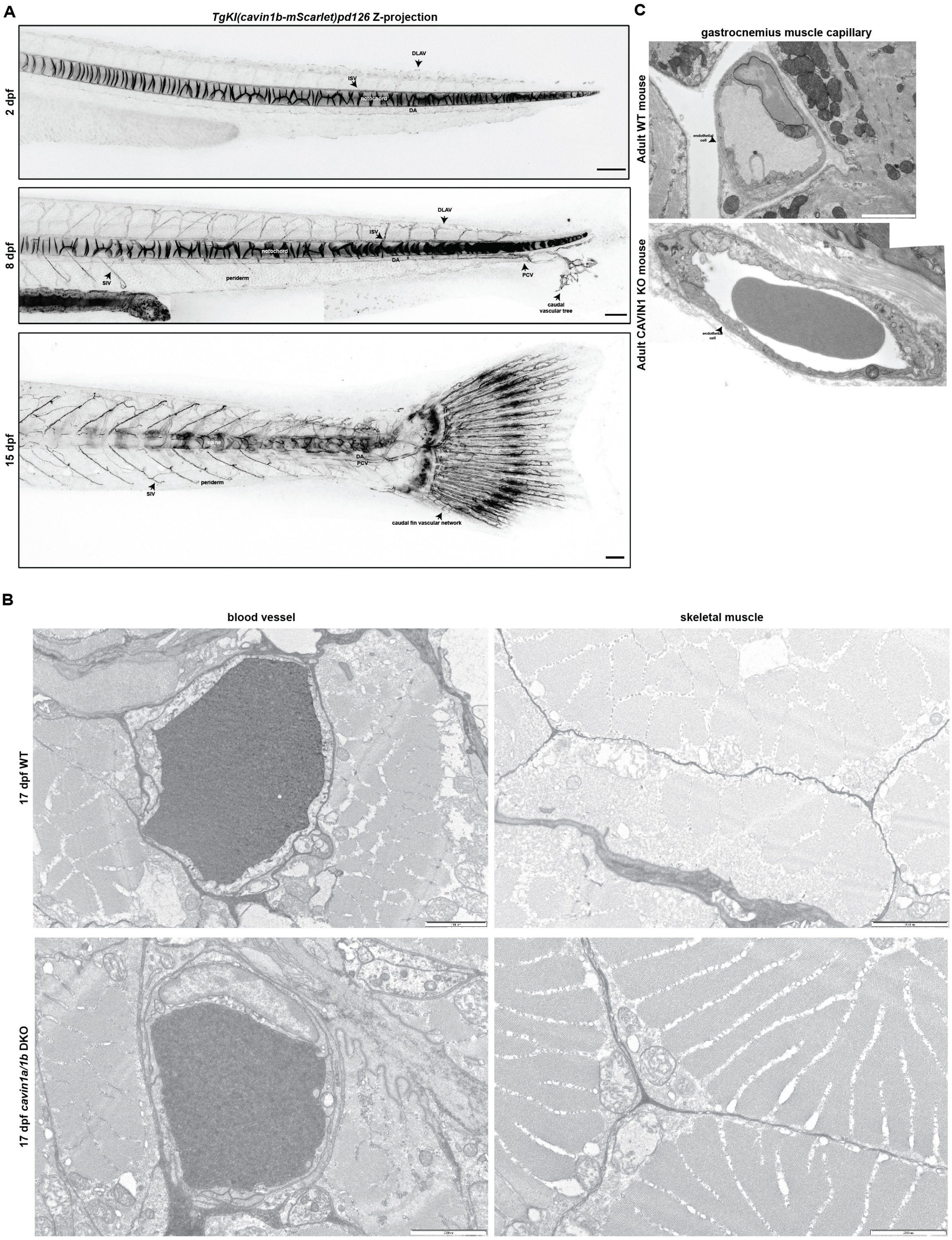


**Figure S5**. **(A)** Tiled confocal images of 2 dpf, 8 dpf and 15 dpf *TgKI(cavin1b-mScarlet)$^{pd126}$* transgenic zebrafish expressing cavin1b-mScarlet. Scale bar: 100 µm. **(B)** Transmission electron micrograph of a gastrocnemius muscle capillary of adult WT and CAVIN1 KO mice. Scale bar: 2 µm **(C)** Transmission electron micrograph of the blood vessels and skeletal muscles of 16 *dpf WT and icavin1a/1b DKO* zebrafish lines. Scale bar: 2 µm ISV=intersegmental vessel; DA=dorsal aorta; DLAV=dorsal anastomosing vessel; SIV=superficial intersegmental blood vessel; PCV=posterior cardinal vein.

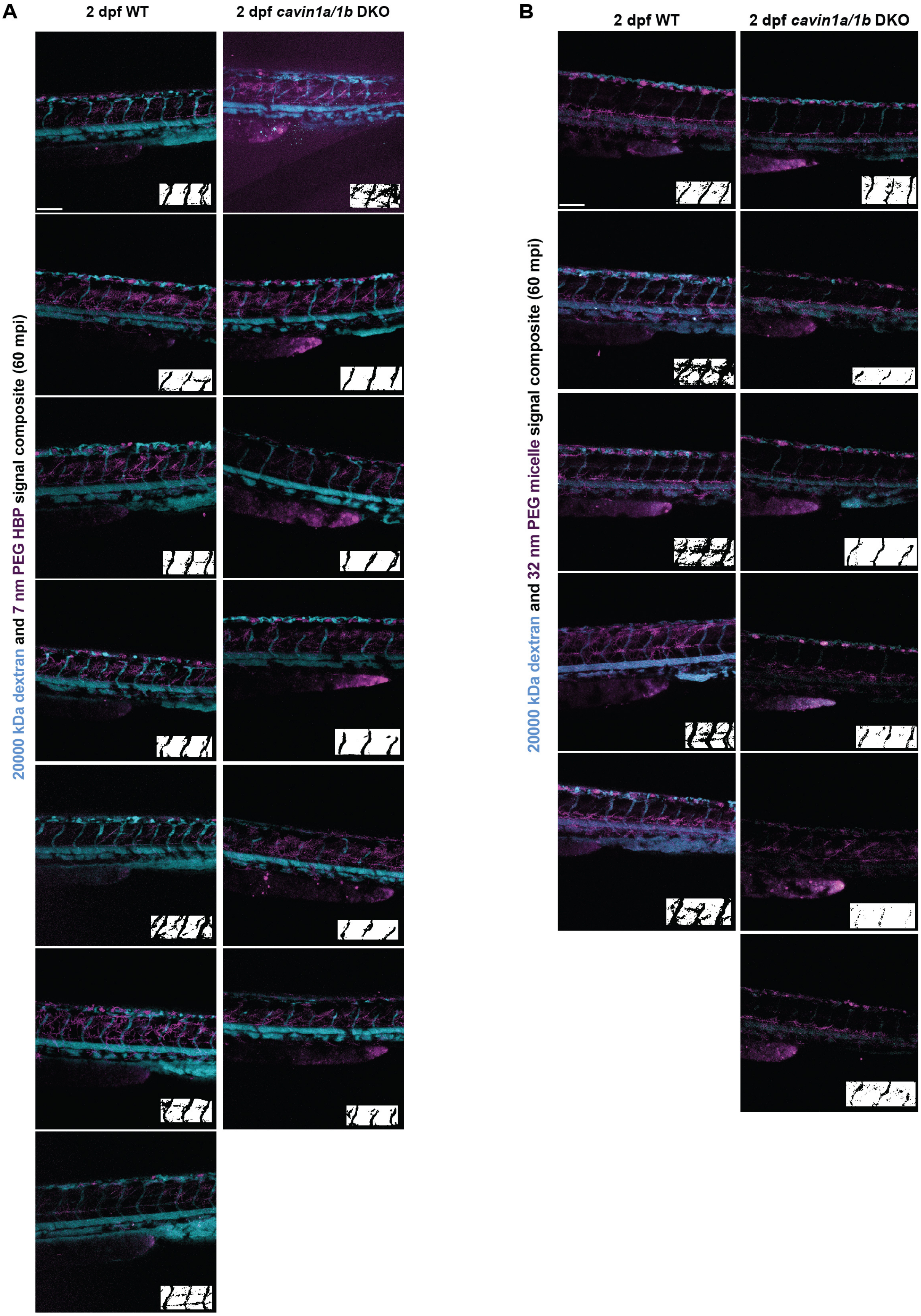

A
2 dpf WT
2 dpf cavin1a/1b DKO
20000 kDa dextran and 7 nm PEG HBP signal composite (60 mpi)
B
2 dpf WT
2 dpf cavin1a/1b DKO
20000 kDa dextran and 32 nm PEG micelle signal composite (60 mpi)

**Figure S6**. **(A)** Z-projected confocal images of each microinjected 2 dpf zebrafish analyzed in **Figure 6C**, with insets showing corresponding threshold masks generated by the signal measurement macro. Scale bar: 100 µm **(B)** Z-projected confocal images of each microinjected 2 dpf zebrafish analyzed in **Figure 6D**, with insets showing corresponding threshold masks generated by the signal measurement macro. Scale bar: 100 µm

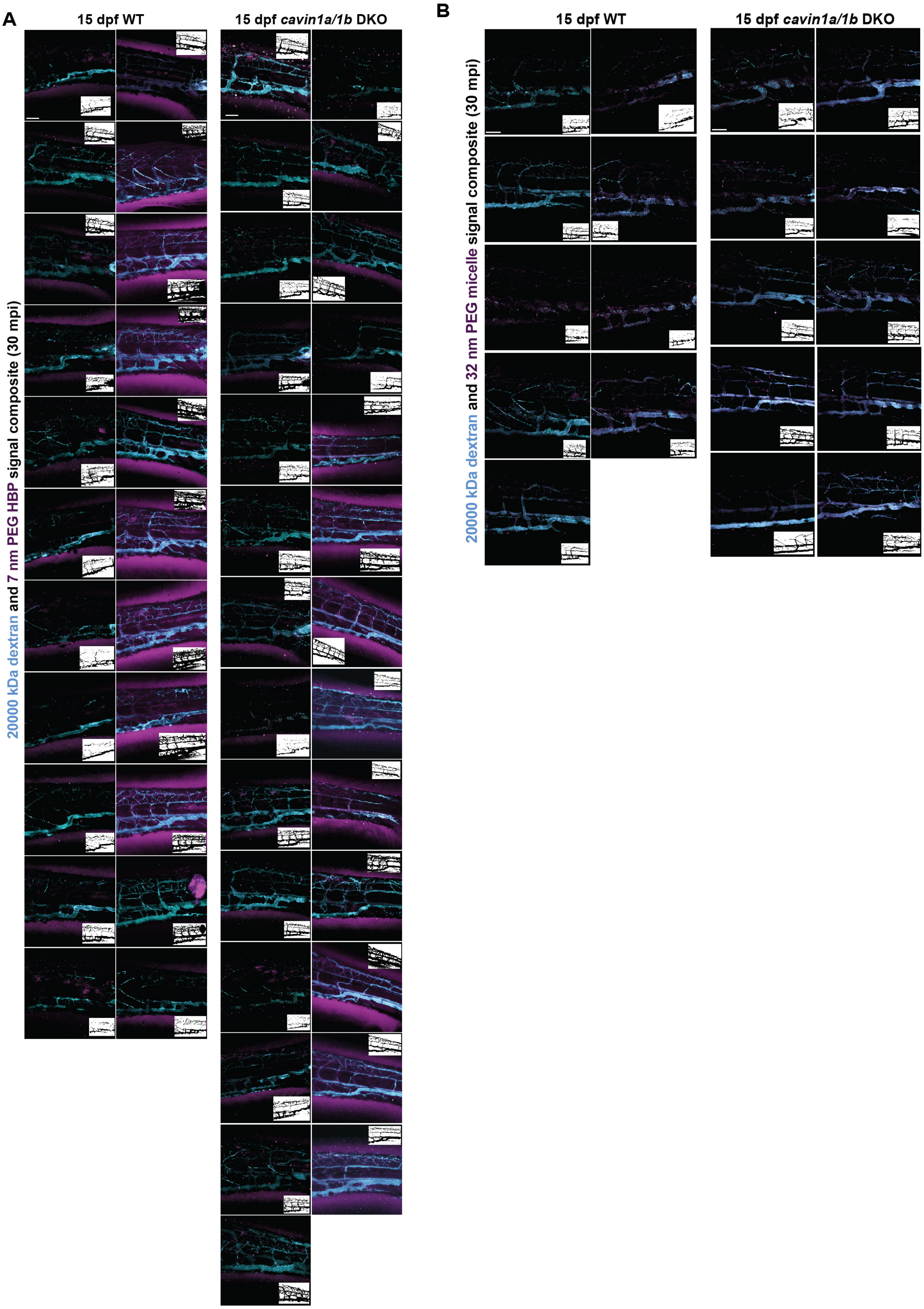


**Figure S7**. **(A)** Z-projected confocal images of each microinjected 15 dpf zebrafish juvenile analyzed in **Figure 6E-F**, with insets showing corresponding threshold masks generated by the signal measurement macro. Scale bar: 100 µm **(B)** Z-projected confocal images of each microinjected 15 dpf zebrafish juvenile analyzed in **Figure 6G-H**, with insets showing corresponding threshold masks generated by the signal measurement macro. Scale bar: 100 µm
